# Armed with PRICKLE(3)s: Stabilizing WNT/PCP complexes against RNF43-mediated ubiquitination

**DOI:** 10.1101/2025.03.24.644882

**Authors:** Katarzyna A. Radaszkiewicz, Tomasz W. Radaszkiewicz, Pavla Kolářová, Petra Paclíková, Kristína Gömöryová, Tomáš Bárta, Kateřina Hanáková, Zbyněk Zdráhal, Jakub Harnoš

## Abstract

The human Prickle protein family, consisting of PRICKLE1, PRICKLE2, PRICKLE3, and PRICKLE4, is an integral component of the WNT/planar cell polarity (WNT/PCP) pathway and is essential for various cellular and developmental processes. Despite their significance, the detailed roles and involvement in molecular mechanisms of these proteins in cells remain not fully understood. In this study, we used enhanced proximity biotinylation (miniTurboID) combined with mass spectrometry to characterize the microenvironment of PRICKLE1-3. Our results reveal that PRICKLE3 is directly linked to the WNT/PCP pathway, primarily localizing at the plasma membrane and forming complexes with VANGL proteins. This observation prompted us to examine its role in the non-canonical WNT signalling pathway in more detail. Using an inducible expression system to achieve protein levels closer to physiological conditions, we found that PRICKLE3 enhances the stability of VANGL1 and VANGL2 by shielding them from Casein kinase 1 ε-mediated phosphorylation. Furthermore, our results indicate that PRICKLE3 modulates WNT receptor complexes by negatively regulating the interaction between Casein kinase 1 ε and ubiquitin ligase RNF43, resulting in decreased ubiquitination and increased stabilization of VANGL1/2 at the plasma membrane. Notably, these effects were specific to PRICKLE3, with PRICKLE1 showing no comparable activity. Contrary to previous findings based mainly on standard overexpression studies, neither PRICKLE3 nor PRICKLE1 influenced the levels or phosphorylation status of WNT proteins DISHEVELLED2 and DISHEVELLED3, which are the PRICKLE proteins binding partners. In summary, we have identified a key mechanism specific to PRICKLE3 that positively regulates WNT/PCP complexes by suppressing RNF43. Additionally, we present a comprehensive interactome and new tools for the functional specification of Prickle isoforms to support further research.

## INTRODUCTION

The planar cell polarity (PCP) pathway is the most extensively studied branch of non-canonical, β-catenin-independent WNT signaling (1,2). A hallmark of the PCP pathway is its ability to propagate polarized information between neighbouring cells, ultimately coordinating the global orientation of tissues within the body plane (3,4). This foundational concept of PCP signaling was first established through pioneering genetic studies in *Drosophila melanogaster*, where several key PCP genes were identified (4–6). In vertebrates, the PCP pathway not only governs cellular polarity within tissues but also orchestrates essential processes such as collective cell migration, individual cell motility, and axon guidance (6–9). Disruptions in PCP signaling are associated with a variety of developmental defects, including neural tube closure anomalies, such as spina bifida, scoliosis, and Robinow syndrome (10–12). Moreover, this pathway is often dysregulated in cancer, driving metastatic events and resistance to treatment (13,14).

The propagation of PCP signaling depends on antagonistic interactions between specific protein complexes, particularly those formed by Frizzled (FZD) receptors and the cytosolic hub protein Dishevelled (DVL), which compete with the plasma membrane–spanning protein Van Gogh-like (VANGL) and its cytosolic partner, Prickle (PK) (5,6,15). Another critical PCP component is the transmembrane protein Flamingo, known in vertebrates as Cadherin EGF LAG Seven-Pass G-Type Receptor (CELSR) (16). In vertebrate systems, PCP-related WNT receptor complexes include additional co-receptors such as Receptor tyrosine Kinase-like Orphan Receptor 1/2 (ROR1/2), Receptor-like Tyrosine Kinase RYK, and Protein Tyrosine Kinase 7 (5,6). Typical signaling molecules that activate this pathway include the WNT ligands such as WNT5A or WNT11 that activate PCP signaling (1,2,5,6).

Despite the PCP pathway’s broad physiological relevance, the PRICKLE protein family remains understudied, particularly in human cells. In this study, we focus specifically on human PRICKLE1–3 isoforms. While some general insights are available on these proteins, their specific roles within human PCP signaling remain unclear (11,17). Proper establishment of planar polarity depends on various mechanisms, including protein trafficking and stability, post-translational modifications, actin cytoskeleton regulation via small GTPases, actomyosin network contractility, and the expression and propagation of WNT ligands (2,9,14,18,19).

However, how PRICKLE proteins specifically contribute to these mechanisms within the WNT/PCP pathway remains unclear. Do all PRICKLE isoforms act similarly or differently? Are they positive or negative regulators of the WNT5A pathway transduction? What are their intracellular localizations, and how are they regulated?

To address these knowledge gaps, we investigated the roles of individual PRICKLE isoforms in human cells using enhanced proximity-dependent biotinylation coupled with mass spectrometry, allowing us to map PRICKLE interactions at a detailed level. Our findings reveal that PRICKLE3 exhibits a particularly pronounced connection to core WNT/PCP proteins, promoting the stability of VANGL and potentiating the activity of WNT5A receptor complexes. Moreover, we identified the involvement of the PRICKLE3 in the regulation of the clinically relevant E3 ubiquitin ligase – Ring Finger protein 43 (RNF43). These findings show that PRICKLE3 has a unique role in regulating WNT/PCP signaling and that we provide useful tools for studying PRICKLE isoforms in human cells.

## RESULTS

### Proximity interactomes of PRICKLE1, PRICKLE2 and PRICKLE3

Proximity-dependent labelling techniques have emerged as a well-established method for mapping protein-protein interactions (20,21). To characterize the interactome of the selected PRICKLE 1-3 proteins, we employed an enhanced proximity-dependent biotinylation (miniTurboID) method. MiniTurboID technique relies on the action of the modified version of promiscuous biotin ligase BirA*, which can biotinylate both stable and transient interacting proteins in a radius of approximately 10 nm around the protein of interest (POI, bait) (**Fig. 1A**) (22). Importantly, cell lysis can be performed in the presence of strong detergents, which helps to isolate hydrophobic proteins – an advantage over standard IP experiments. Moreover, this pull-down utilizes extraordinarily strong non-covalent interaction between biotin and streptavidin for isolation of the reaction products. This technique provides a powerful tool for mapping protein interactions, especially for hydrophobic or transient interactors (21).

**Figure 1.**
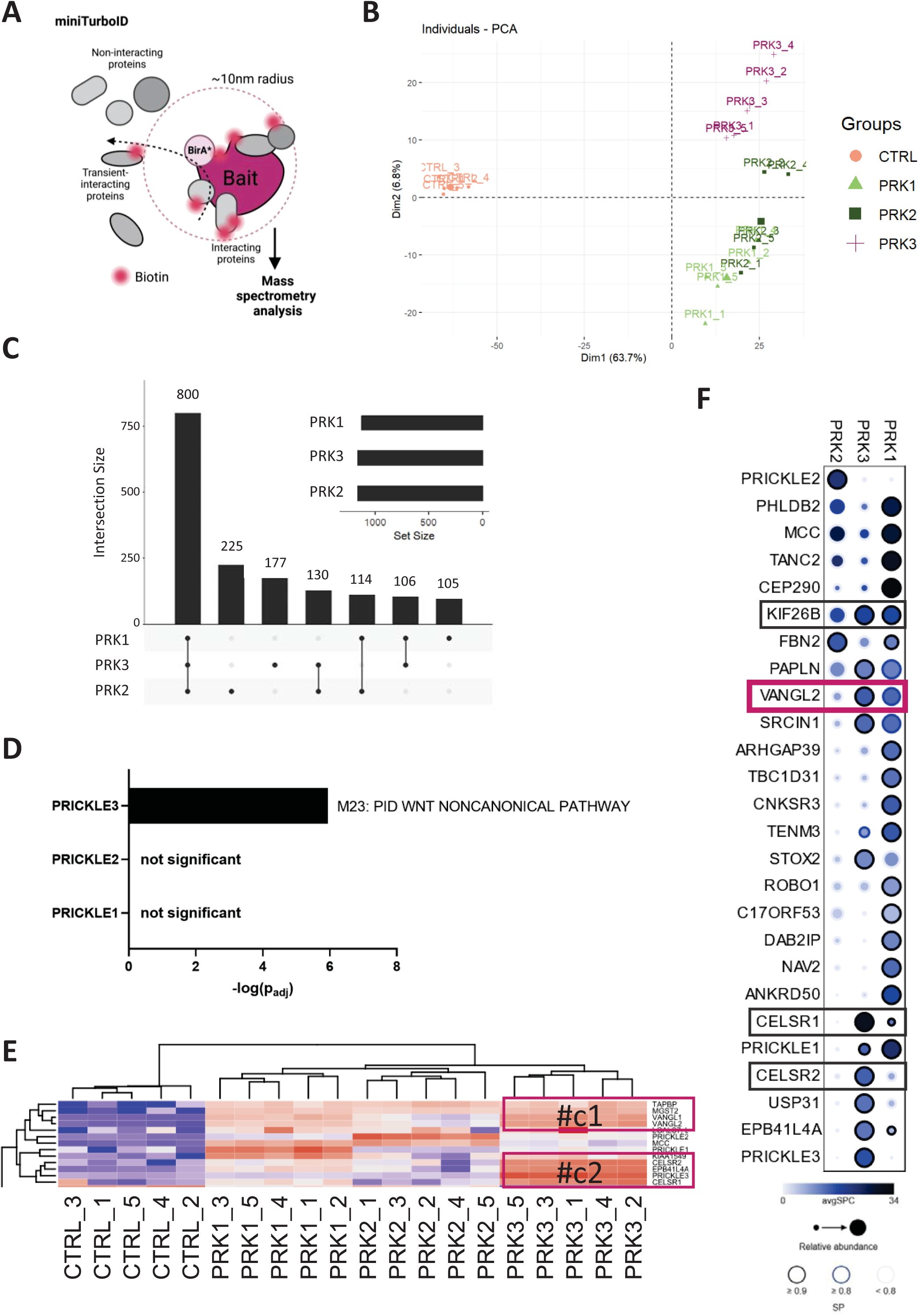
Proximity interactomes of PRICKLE1, PRICKLE2 and PRICKLE3. A) Scheme of the miniTurbo proximity-dependent biotinylation (miniTurboID) assay. Created with BioRender.com B) The Principal Component Analysis (PCA) demonstrates clustering corresponding to specific PRICKLE variants, while the individual replicates for each bait are grouped within their respective clusters. C) The UpSet plot illustrates the overlap of upregulated preys across all PRICKLE (1–3) variants, providing insight into the common and distinct interactions across these proteins. D) The Metascape GO terms analysis reveals that only the interactors of PRICKLE3 were significantly enriched in the components of the Planar Cell Polarity (PCP) pathway. E) Fragment of a heatmap of proteins upregulated for at least one bait. Using k-means clustering, two distinct clusters of preys, labelled as #C1 and #C2, were identified. Both clusters contain preys specific to the Planar Cell Polarity (PCP) pathway. F) The dot plot presents the results of the SaintExpress analysis, which verifies the bait-prey interactions for individual clusters. The colour of the dots indicates the abundance of the preys, with darker shades representing higher abundance. The size of each dot corresponds to the relative abundance of the prey, while the edge colour reflects the Saint Probability, with darker colours indicating stronger bait-prey interactions.

To map PRICKLE interactomes, we developed stable HEK T-REx 293 TetON cell lines that inducibly express the PRICKLE isoforms 1-2-3 fused N-terminally with modified BirA* ligase (miniTurboID) and the V5 tag. Cells expressing only miniTurboID-V5 served as controls (**Supp. Fig. 1A**). We validated the specificity of biotinylation by Westen Blot (WB) and immunofluorescence, confirming that interacting proteins were biotinylated only in the presence of doxycycline (DOX) and biotin (**Supp. Fig. 1B-E**). We prepared five biological replicates for each POI. The biotinylated proteins were enriched using streptavidin-coated beads and subsequently identified through mass spectrometry (MS). In total, MS identified a set of 7011 proteins, which after the quality filtering was reduced to 5986 proteins (**Suppl. Table 1**). Subsequently, principal component analysis (PCA) of the 500 most variable proteins (**Fig. 1B**) revealed cohesive clustering of replicates and excluded the batch effect. Importantly, it shows that PRICKLE3 has the most distinct interactome, whereas PRICKLE1 and PRICKLE2 share similar baits (**Fig. 1B**).

All PRICKLE variants had a similar number of significant proximal interactors, around 1100 of preys (**Supp. Fig. 2A**, **Suppl. Table 2**), sharing approximately 800 (**Fig. 1C**). PRICKLE2 had the highest number of unique interactors (225), followed by PRICKLE3 (117) and PRICKLE1 (105). Interestingly, PRICKLE3 and PRICKLE2 shared most of the candidate interactors (130), while PRICKLE3 and PRICKLE1 shared 106 preys (**Fig. 1C**, **Suppl. Table 3**). This implies a high level of redundancy between PRICKLE1- 3 together with some degree of specialization of the individual POIs. Next, Gene Ontology (GO) analysis (23) showed that the interactome of PRICKLE1-3 is enriched with proteins associated with organisation of cell membrane and cytoskeleton, as well as regulation of metabolism and other processes (**Suppl. Fig. 2D**). Moreover, GO analysis revealed that all PRICKLEs are involved in various signalling pathways, especially signalling by Vascular endothelial growth factor (VEGF), Rho GTPases and WNT signalling pathway (**Suppl. Fig. 2D**). Subsequently, we used the Human Cell Map (HCM) (24) to precisely determine the cellular localisations of PRICKLE1-3. A great advantage of using this resource is that it was developed with the BioID assay in the same cellular model as our study (i.e., T-REx 293 cells). Our analysis indicated that PRICKLE1 and PRICKLE3 were predominantly located in cell junctions whereas PRICKLE2 was surprisingly associated with the mitochondrial matrix (**Suppl. Fig. 2B**, **Suppl. Table 4**). However, not all proteins identified in this study could be mapped to the HCM, likely due to the use of a more robust biotin ligase – miniTurboID.

In this research, we focused on the WNT signaling pathway due to its significant role in disease and developmental processes (6,14,18). PRICKLE proteins are primarily associated with this pathway, acting as key regulators of neurulation, body axis elongation, and organogenesis (11). Disruptions in the balance or mutations of PRICKLE proteins have been linked to developmental abnormalities, including neural tube defects, as well as to cancer progression and various neurological and neurodegenerative disorders (11). However, the mechanism of their action is not described well, and little is known about the functions of individual isoforms. Thus, the GO analysis of the interactors specific to the individual PRICKLE proteins helped to determine their potential individual roles in the WNT signalling. Indeed, we observed that PRICKLE3 is connected the most with the noncanonical WNT signalling pathway (**Fig. 1D**, **Suppl. Table 5-8**). Furthermore, *k*-means clustering of all candidate interactors (**Fig. 1E**, **Suppl. Fig. 2C**) uncovered two clusters of PRICKLE3 interactors, composed of known protein families involved in the WNT/PCP signalling – VANGL1/2 and CELSR1/2 (16,18).

As a complementary approach to determining the confident bait-prey interactions using logFC and adjusted p-values threshold on limma test results, we employed the SAINTexpress tool (25) integrated within the REPRINT resource (26). SAINTexpress provides the SAINT probability (SP) metric, where SP>0.9 means a highly confident interaction. In our data, we identified 26 bait-prey pairs visualized by DotPlot using SP>0.9 (**Fig. 1F**, **Suppl. Table 9**). This analysis further supports that PRICKLE3 isoform is the most significantly related to the components such as VANGL2, CELSR2, and Kinesin-like protein KIF26B (**Fig. 1F**) (16,18,27). These findings reinforce PRICKLE3’s specialized role within the WNT/PCP signaling pathway.

This study provides a rich and validated resource of the PRICKLE1-3 interactomes for further investigation. For subsequent mechanistic studies, we selected PRICKLE3 as a poorly described isoform and strongly associated with the noncanonical WNT pathway which is crucial for development and disease (11,17,28). This isoform was then compared with PRICKLE1 to determine whether it plays a distinct role in the WNT signaling cascade. Based on the HCM data, both of these PRICKLEs are localised at the cellular periphery where WNT ligands reception occurs. However, a more detailed analysis, involving the plotting of data as log fold change (log FC) against -log10 (adjusted p-value, padj), reveals further insights (**Suppl. Fig. 2E**). This approach suggests that a bait protein is likely to remain longer and result in stronger biotinylation of prey proteins, or alternatively, indicates a higher concentration of the bait protein within a particular cellular compartment. Through this method, it becomes evident that PRICKLE3 exhibits a stronger association with integral plasma membrane WNT components than PRICKLE1, as demonstrated by: co-receptor low-density lipoprotein receptor-related protein 6 (LRP6), FZD3, ROR1/2, CELSR1/2. Moreover, the applied method also indicates a stronger association of PRICKLE3 with VANGL proteins, consistent with previous findings (29,30). Contrary, PRICKLE2 is localised predominantly in mitochondria. Thus, we did not follow it in the next parts of our work. This resource sets a foundation for further research, with PRICKLE3’s unique role in WNT signaling at the membrane underscoring its importance.

### PRICKLE3 is a positive regulator of the WNT/PCP pathway

All PRICKLE (1–3) protein family isoforms (**Fig. 2A**) share three sequence elements: the N-terminal PET (Prickle, Espinas, Testin) domain, involved in protein-protein interactions and signal transduction; several LIM (Linl-1, Isl-1, Mec-3) domains, which are cysteine-rich protein modules with diverse functions; and the C-terminal PKH (Prickle homology) domain with less determined role. Furthermore, all PRICKLE proteins contain a central intrinsically disordered region that varies among isoforms. Notably, unlike PRICKLE1 and PRICKLE2, PRICKLE3 lacks the Vangl binding motif (VBM) involved in binding to the intracellular region of VANGL (11). Based on this fact and our proteomic data, this variation in VBM in PRICKLE3 suggests it may operate through an alternative mechanism in cells.

**Figure 2.**
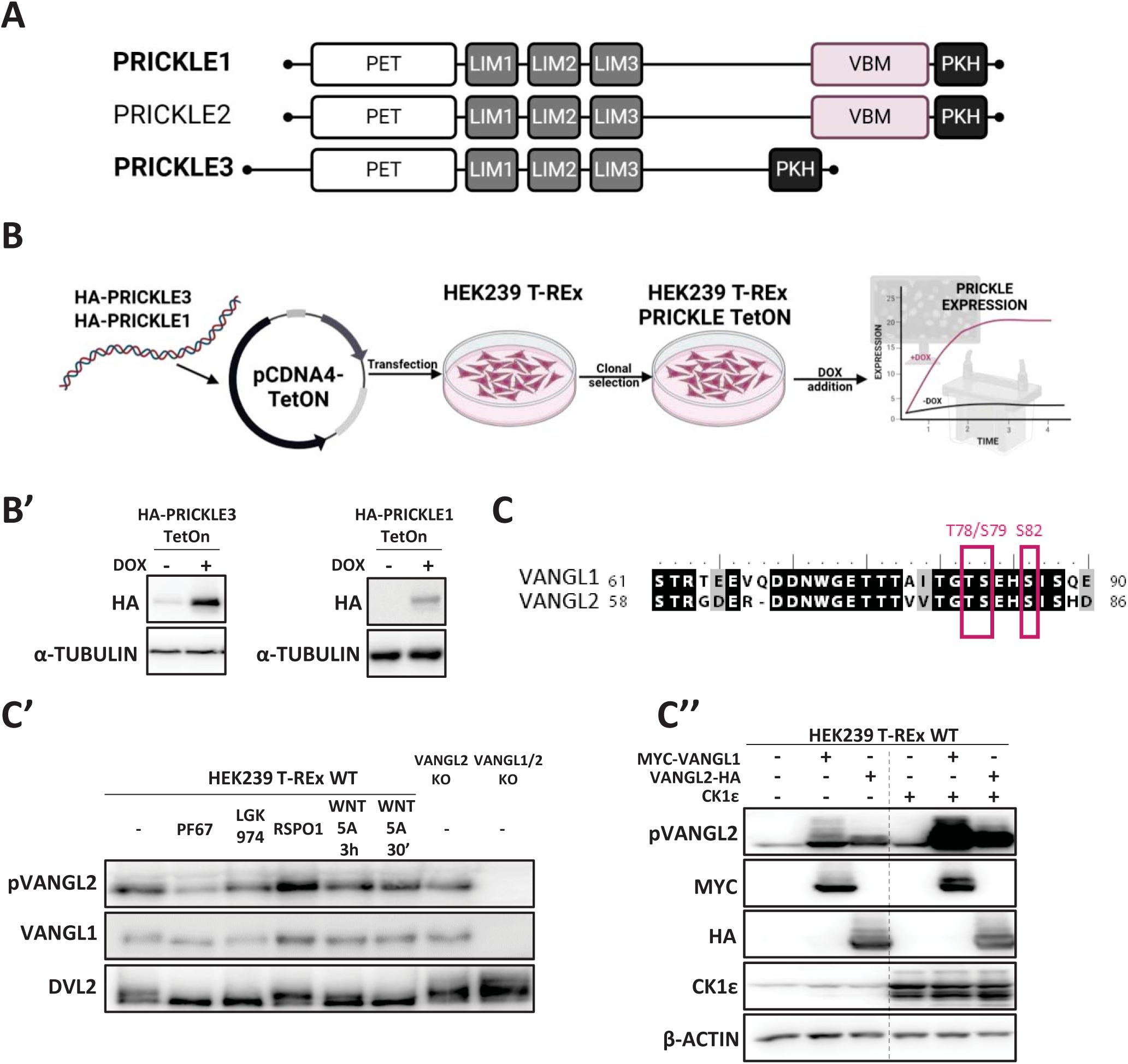
VANGL1/2 research toolbox. A) PRICKLE1-3 sequence elements. For a complete list of descriptions and abbreviations, please refer to the accompanying text. Created with BioRender.com B) Preparation of stable and clonal doxycycline-inducible cell lines expressing N-terminally HA-tagged PRICKLE1 and PRICKLE3. The HEK T-REx 293 wild type (WT) cell line was transfected with selected constructs, and single clones were subsequently derived and validated. B’) The expression of PRICKLE1/3 was validated in the presence of doxycycline through Western blotting, confirming the successful induction of PRICKLE expression in the cell lines. α-TUBULIN served as loading control. Created with BioRender.com C) CK1ε phosphorylation sites of VANGL1/2 protein. C’) Validation of a-Vangl antibodies at the endogenous level. HEK T-REx 293 WT cells were pre-treated over night with porcupine inhibitor LGK- 974 to block Wnt ligands secretion and subsequently stimulated with 100 ng/ml of human recombinant WNT5A for 3 hours or 30 min; 25ng/ml of R-Spondin1 (RSPO1) or inhibitor of CK1ε/δ - 2.5 μM PF-670462 (PF67) for 3h. HEK T-REx 293 VANGL2 knockout (KO) and VANGL1/2 double knockout (KO) cells were utilized as controls to assess antibody specificity. C’’) Validation of Vangl antibodies specificity – overexpression model. HEK T-REx 293 wild type (WT) cells were transfected with plasmids encoding MYC-tagged VANGL1, HA-tagged VANGL2, and CK1ε and analysed by Western blotting.

For functional studies, we generated stable HEK T-REx 293 TetON cell lines that inducibly express PRICKLE1 and PRICKLE3 isoforms fused N-terminally with the HA tag upon doxycycline treatment (**Fig. 2B,B’**; **Supp. Fig. 3A-A’,B-B’**). The HEK T-REx 293 cell line serves as a well-established model for investigating components and molecular mechanisms of canonical and non-canonical WNT pathways (31,32). Since the VANGL1/2 protein emerged as one of the key preys in our PRICKLE3 protein interaction analysis, we proceeded to validate a set of antibodies that were employed as readouts in subsequent experiments. The N-terminal region of VANGL contains a cluster of serine/threonine residues (T78, S79, and S82 in human VANGL2), which are phosphorylation targets for Casein kinase I epsilon and delta (CK1ε/δ) (**Fig. 2C**). Phosphorylation of CK1ε/δ is required for VANGL activation and its transport to the plasma membrane (33–35). We confirmed that phosphorylation of VANGL1/2 is CK1ε-dependent in our model, as treatment of cells with the CK1ε/δ inhibitor PF-670462 (36) decreased phosphorylation of endogenous VANGL2 and enhanced VANGL1 electrophoretic migration on SDS-PAGE/WB (**Fig. 2C’**). In agreement with that, the overexpression of CK1ε further increased phosphorylation of both VANGL1 and VANGL2 (**Fig. 1C’’**). In our model, the phosphorylation of VANGL was not dependent on the WNT ligands presence as treatment of cells with the WNT secretion inhibitor LGK-974, as well as the WNT5A ligand, did not alter its phosphorylation level (**Fig. 1C’**). Additionally, we assessed antibody specificity in VANGL2 knockout (KO) and VANGL1/2 double KO cells (**Fig. 1C’**) (37). The pVANGL2 antibody used in our study recognizes both isoforms of VANGL; therefore, it was considered a reliable readout for detecting both isoforms (VANGL1/2 or VANGL) in our subsequent analyses.

Previous studies have demonstrated a mutual regulatory relationship between Prickle and Vangl proteins, with evidence suggesting that both can modulate the other’s levels as well influence other components of the WNT pathway (38–44). Considering this, our initial experiment aimed to investigate how PRICKLE3 affects the levels of VANGL and its phosphorylation. Forced expression of PRICKLE3 increased the level of VANGL1 and enhanced its migration on SDS-PAGE, indicating a lower phosphorylation level. This finding was further supported by the pVANGL2 antibody analysis (**Fig. 3A,A’**). To assess the proportion of the CK1ε/δ-activated form of VANGL, we calculated the ratio of phosphorylated VANGL2 to total VANGL1 level, which also showed a decrease (**Fig. 3A’**). Interestingly, contrary to previous reports based on standard overexpression (41,42,45), PRICKLE3 did not affect the phosphorylation of DVL2/3 or its protein level in our system (**Fig. 3A**).

**Figure 3.**
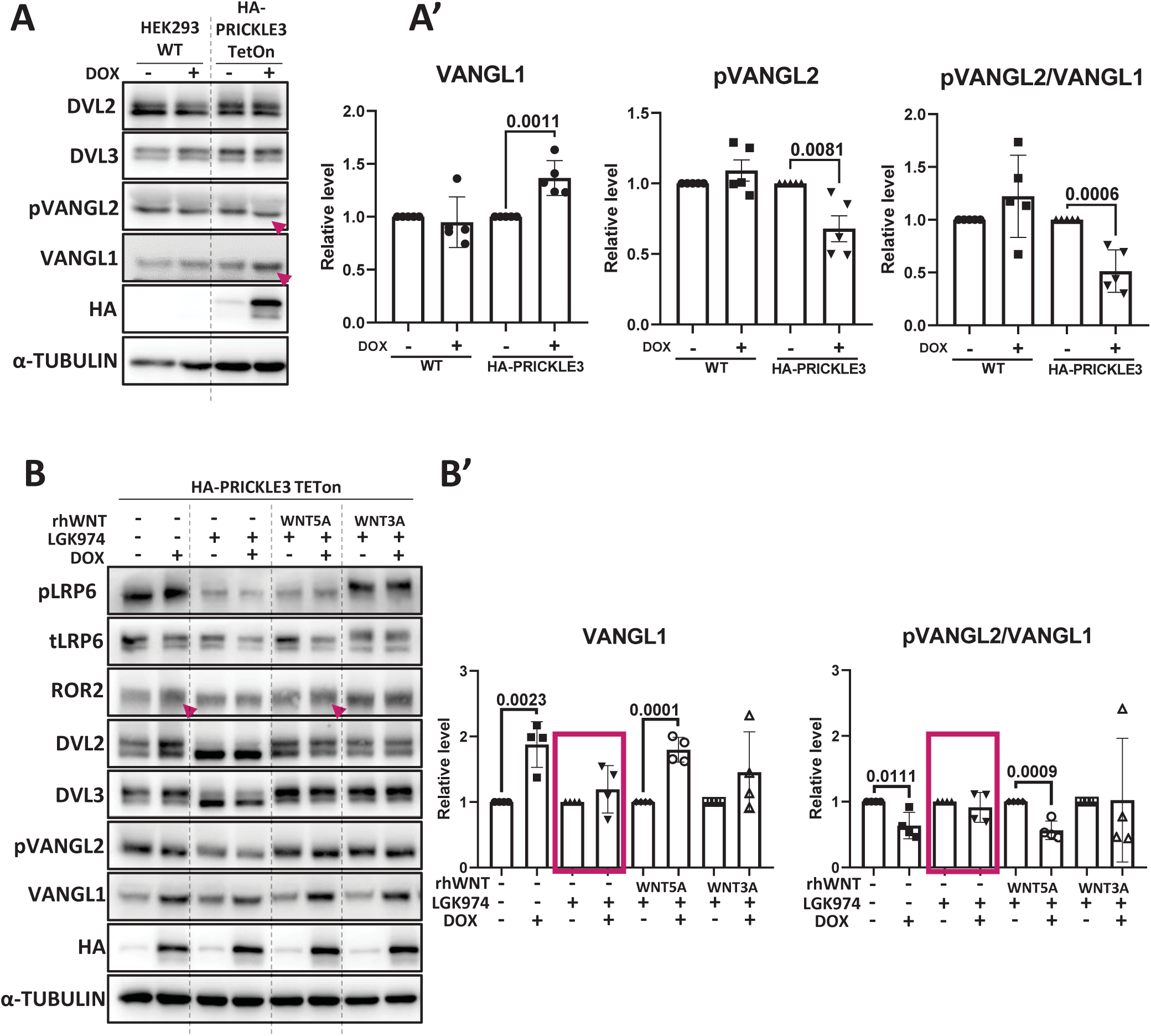
Prickle3 supports active WNT5A receptor complexes. A) Doxycycline-induced overexpression of the PRICKLE3 protein in HEK T-REx 293 PRICKLE3 TetON cell line (HA-PRICKLE3). HEK T-REx 293 wild-type (WT) cells served as a control for the effects of doxycycline. Arrowheads show phosphorylation-dependent change in the electrophoretic mobility of VANGL. α-TUBULIN served as loading control. A representative experiment from N = 4. A’) Densitometric quantification of Western blot (WB) band intensities. Results were normalized to doxycycline-untreated cells. Statistical analysis was performed using an unpaired t-test, and the corresponding *p*-values are reported. N = 4. B) Effect of recombinant WNT treatment. HEK T-REx 293 PRICKLE3 TetON cells were pre-treated over night with porcupine inhibitor LGK-974 to block endogenous Wnt ligands secretion and subsequently stimulated with 100 ng/ml of human recombinant WNT5A or WNT3A for 3 hours. Arrowheads show phosphorylation-dependent change of the electrophoretic mobility of VANGL; α-TUBULIN served as loading control. A representative experiment from N = 4. B’) Densitometric quantification of Western blot (WB) band intensities. Results were normalized to doxycycline-untreated cells. Statistical analysis was performed using an unpaired t-test, and the corresponding *p*-values were reported. N = 4.

Next, we analysed VANGL levels and its CK1ε/δ-mediated phosphorylation status in the presence of LGK-974, as well as under treatment with recombinant WNT5A or WNT3A ligands (**Fig. 3B, B’**). PRICKLE3 had no effect on VANGL1 and pVANGL2/VANGL1 levels in the WNT ligands absence, suggesting that it affects only active WNT/PCP receptor complexes. Consistently, the phenotype related to the PRICKLE3 was evident only upon non-canonical WNT5A ligand treatment, but not the WNT3A involved in canonical WNT signalling. Moreover, PRICKLE3 overexpression led to a shift in the electrophoretic motility of ROR2 (see arrowhead). This protein acts as the WNT5A co-receptor and it is phosphorylated upon activation by WNT ligand binding (46,47). It was also shown that ROR2 binds VANGL2 in response to WNT5A, forming a receptor complex of the WNT/PCP pathway (34). This observation supports the role of PRICKLE3 in the regulation of the non-canonical WNT pathway via involvement in the active WNT/PCP complexes at the plasma membrane. Similarly to our previous results, PRICKLE3 did not influence the level and phosphorylation status of DVL2/3 (**Fig. 3B**).

The increased levels of VANGL protein in the presence of PRICKLE3 suggest that PRICKLE3 may either protect VANGL from degradation or promote its synthesis. To validate the interaction between PRICKLE3 and VANGL, we performed an immunoprecipitation assay, which confirmed that endogenous VANGL, both in non-phosphorylated and phosphorylated forms, interacts with PRICKLE3 (**Fig. 4A**). We then employed a widely used approach to study protein degradation, utilizing the protein translation inhibitor cycloheximide (CHX) (48) in a pulse-chase assay (**Fig. 4B-B’’**). We found that in the presence of PRICKLE3, the level of VANGL is stabilized over time (**Fig. 4B,B’**), confirming that PRICKLE3 protects VANGL from degradation. However, PRICKLE3 had no effect on the VANGL pool phosphorylated by CK1δ/ε (**Fig. 4B,B’’**), and there was no significant effect on DVL2 (**Fig. 4B**).

**Figure 4.**
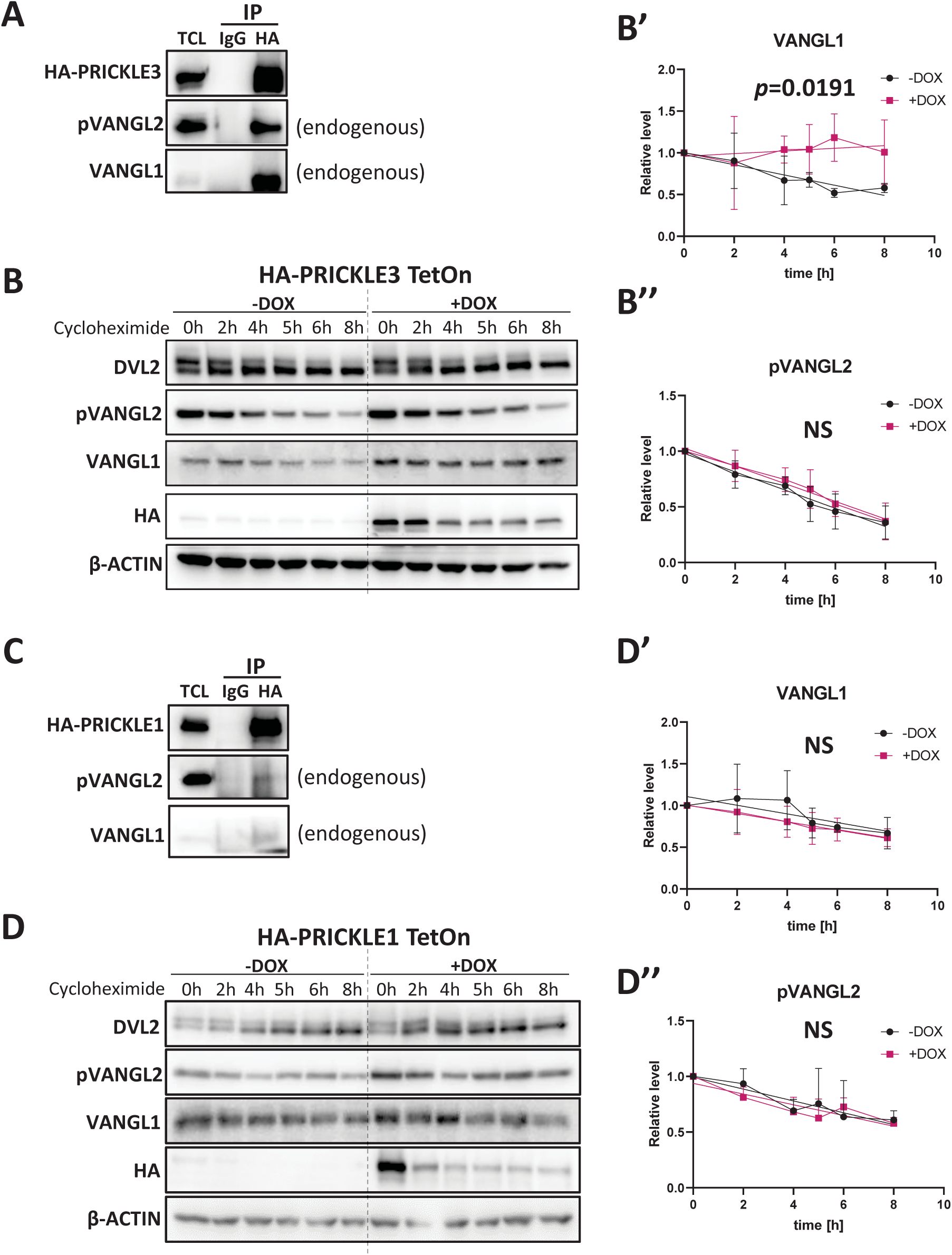
PRICKLE3 not PRICKLE1 protects VANGL2 from degradation. A) Immunoprecipitation of HA-tagged PRICKLE3. HEK T-REx 293 PRICKLE3 TetON cells were induced by ON treatment with doxycycline. Pull-downs using anti-HA tag antibody and IgG control were performed to confirm the interaction between PRICKLE3 and endogenous VANGL1/2. N = 4. B) Protein stability determination using the cycloheximide pulse-chase assay. HEK T-REx 293 PRICKLE3 TetON cells (HA-PRICKLE3) were induced by ON treatment with doxycycline. Subsequently, cells were treated with 50 µg/mL cycloheximide at various time points and analyzed by Western blot. β-ACTIN served as loading control. N = 3. B’,B’’) Densitometric quantification of Western blot (WB) band intensities. Results were normalized to the 0-hour time point. Statistical analysis was performed using linear regression, and the corresponding *p*-values are reported. (NS) indicates no statistical difference. N = 3. C) Immunoprecipitation of HA-tagged PRICKLE1. HEK T-REx 293 PRICKLE1 TetON cells were induced by ON treatment with doxycycline. Pull-downs using anti-HA tag antibody and IgG control were performed to confirm the interaction between PRICKLE1 and endogenous VANGL1/2. N = 4. D) The cycloheximide pulse-chase assay. HEK T-REx 293 PRICKLE1 TetON cells (HA-PRICKLE1) were induced by ON treatment with doxycycline. Subsequently, cells were treated with 50 µg/mL cycloheximide at various time points and analyzed by Western blot. β-ACTIN served as loading control. N = 4. D’,D’’) Densitometric quantification of Western blot (WB) band intensities. Results were normalized to the 0-hour time point. Statistical analysis was performed using linear regression, (NS) indicates no statistical difference, N = 4.

As the next step, we compared the results obtained for PRICKLE3 with PRICKLE1 isoforms. Like PRICKLE3, immunoprecipitation of PRICKLE1 resulted in the pulldown of VANGL and its phosphorylated form (**Fig. 4C**). However, our results indicated that PRICKLE1 did not stabilize VANGL nor its phosphorylated form (**Fig. 4C-C’’**). This is another difference between these two related proteins, as PRICKLE1, unlike PRICKLE3, did not stabilize VANGL protein or promote the active WNT/PCP complexes formation at the plasma membrane. This prompted further examination of the PRICKLE3 mechanism of action in stabilizing VANGL.

### PRICKLE3 protects VANGL from RNF43-mediated ubiquitination by RNF43-CK1ε interaction disruption

Eukaryotic cells employ two major protein degradation mechanisms: the ubiquitin-proteasome system and the autophagy-lysosome pathway (49). Consequently, we concentrated next on the E3 Ubiquitin ligase – RNF43, a transmembrane enzyme that facilitates the removal of the WNT ligands receptors FZDs and LRP5/6 from the cells surface. It can also interact and suppress the core receptor complex components involved in the non-canonical WNT5A-dependent pathway, including ROR1/2, VANGL1/2, DVL1-3, and CK1 kinases (31,50–53), Importantly, RNF43 has been shown to trigger VANGL2 ubiquitination and its proteasomal degradation (31), which aligns with our results.

To investigate the role of PRICKLE3 in RNF43-dependent degradation of VANGL, we performed co-immunoprecipitation experiments, which demonstrated physical interaction between these proteins (**Fig. 5A**). In the next step, we utilized immunofluorescence to confirm that PRICKLE3, VANGL1, and RNF43 colocalize both intracellularly and at the plasma membrane (**Fig. 5B**, see yellow arrowheads). Next, His-ubiquitin pulldown assay was utilized to show RNF43-mediated ubiquitination of VANGL in the presence and absence of PRICKLE3 (**Fig. 5C**, **Suppl. Fig. 4A**). Therefore, VANGL2 was co-expressed with either RNF43 or its enzymatically inactive RNF43 Mut1 variant (31,53), which served as a negative control. As a result, we demonstrated that the ubiquitination of VANGL2 by active RNF43 was greatly reduced in the presence of PRICKLE3. This indicates that PRICKLE3 suppresses the enzymatic activity of the RNF43, leading to the VANGL stabilization.

**Figure 5.**
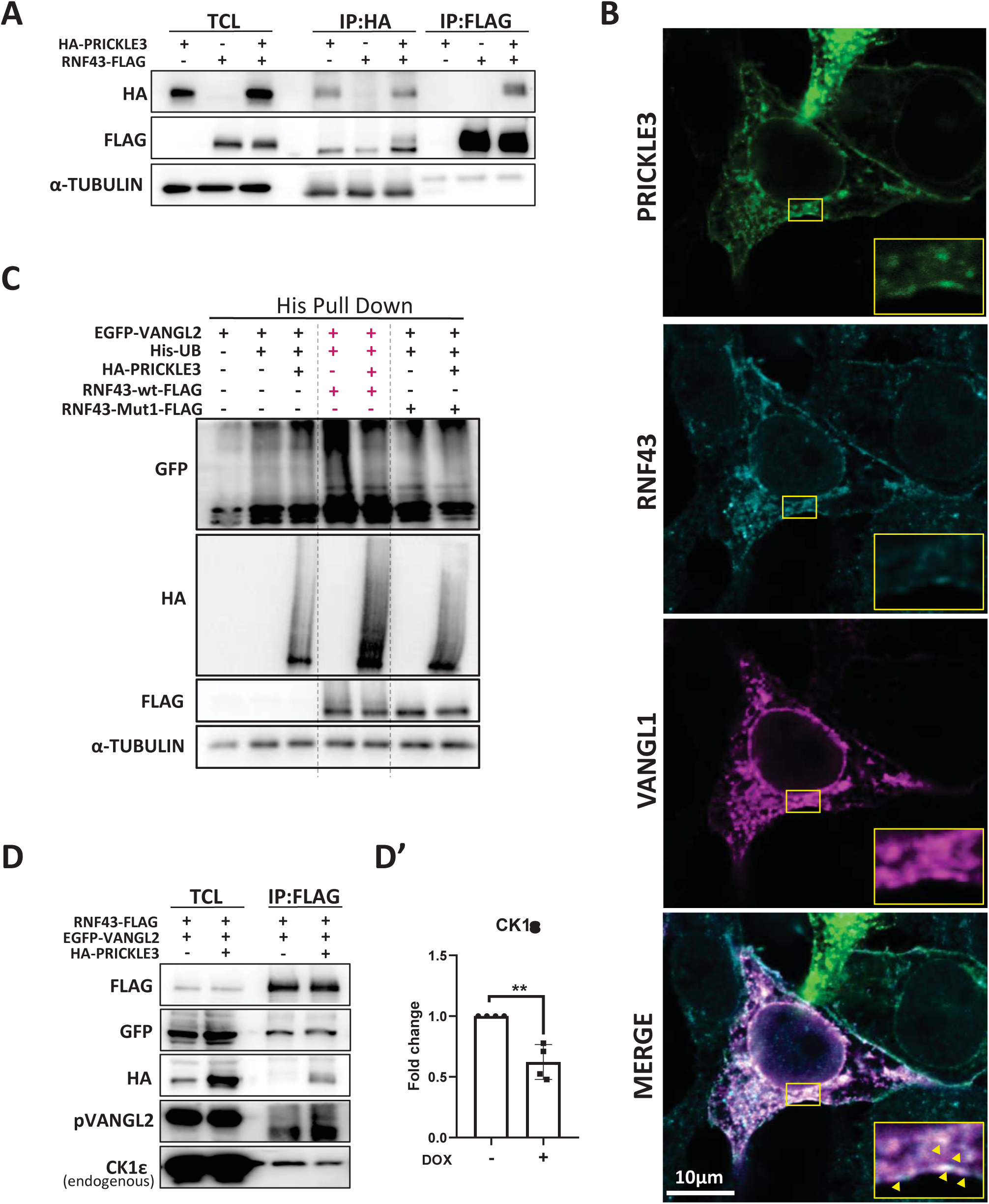
PRICKLE3 protects VANGL2 from RNF43-mediated ubiquitination and proteasomal degradation by CK1e sequestering. A) RNF43 interacts with PRICKLE3. HA-PRICKLE3 was overexpressed with RNF43-FLAG in HEK 293 wild-type cells. Pull-down was made by anti-HA or anti-FLAG antibodies and analyzed by western blotting. TCL- total cell lysate. A representative experiment from N = 3. B) Immunofluorescence imaging demonstrates the colocalization of PRICKLE3, RNF43, and VANGL1. HEK 239 wild-type cells were transfected with HA-PRICKLE3, MYC-VANGL1, and RNF43-FLAG constructs. For protein visualisation anti-tag antibodies were used. Arrowheads indicate points of protein co-localization. C) Ubiquitination assay. HEK T-REx 293 PRICKLE3 TetON cells were transfected His-tagged ubiquitin, EGFP-VANGL2 and FLAG-tagged wild-type or enzymatically inactive Mut1 RNF43 constructs. PRICKLE3 expression was induced by ON treatment with doxycycline. Ubiquitinated proteins were enriched by His pull down and analysed by WB. Representative experiment from N = 3. D) Immunoprecipitation of RNF43-VANGL2 complexes in presence or absence of PRICKLE3 protein. HEK T-REx 293 PRICKLE3 TetON cells were transfected with RNF43-FLAG and EGFP-VANGL2 constructs. PRICKLE3 expression was induced by ON treatment with doxycycline. A representative experiment from N = 4. D’) Densitometric quantification of endogenous CK1ε band intensities. Results were normalized to doxycycline-untreated cells. Statistical analysis was performed using an unpaired t-test, and the corresponding *p*-values are reported. N = 4.

Given the fact that RNF43 activity is regulated by phosphorylation via CK1 family members (50,51), we examined the levels of CK1ε within the VANGL2–RNF43 complexes in the presence of PRICKLE3 (**Fig. 5D**). Overexpression of PRICKLE3 lead to the significant decrease of the CK1ε interacting with RNF43 (**Fig. 5D’**), suggesting that PRICKLE3 may inhibit RNF43 function by interfering with the binding of CK1ε to RNF43.

The importance of Vangl phosphorylation in establishing PCP has been confirmed in a range of animal models, including *Drosophila*, *Xenopus*, *Danio rerio*, and mouse (33–35,44,54). Phosphorylation of Vangl by CK1ε/δ is essential for its activation, prevention of degradation via endoplasmic-reticulum-associated protein degradation pathway, and subsequent transport to the plasma membrane (33). Interestingly, our data demonstrate that PRICKLE3 presence leads to a decreased level of phosphorylated VANGL. Moreover, this form is not stabilized by PRICKLE3 in the CHX pulse-chase experiment. Therefore, we started to investigate the relationship between PRICKLE3, CK1ε, and VANGL proteins. Firstly, immunoprecipitation of PRICKLE3 confirmed its interaction with the endogenous CK1ε (**Fig. 6A**). Next, to evaluate the effect of PRICKLE3 on VANGL phosphorylation by CK1ε, cells were treated with the CK1δ/ε specific inhibitor PF-670462 over a time course (**Fig. 6B,B’**, **Suppl. Fig. 5A**). In the presence of PRICKLE3, the level of phosphorylated VANGL (pVANGL2) did not increase over time, suggesting that PRICKLE3 plays a direct role in preventing VANGL phosphorylation by CK1ε (**Fig. 6B’**).

**Figure 6.**
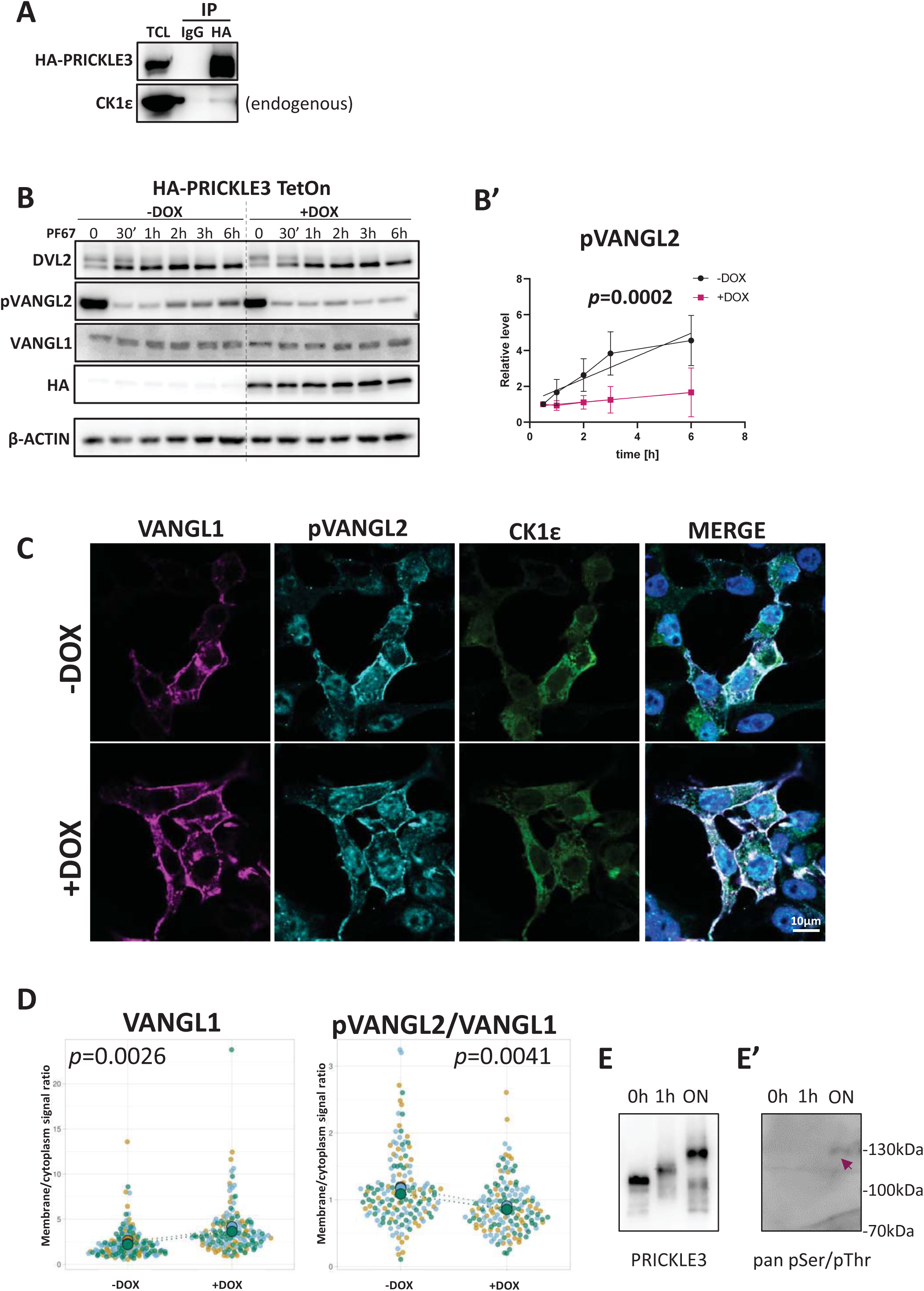
PRICKLE3 protects VANGL1/2 from CK1ε-mediated phosphorylation at the plasma membrane. A) Immunoprecipitation of HA-tagged PRICKLE3. HEK T-REx 293 PRICKLE3 TetON cells were induced by ON treatment with doxycycline. Pull-downs using anti-HA tag antibody and IgG control were performed to confirm the interaction between PRICKLE3 and endogenous CK1ε. N = 4. B) Analysis of VANGL phosphorylation by CK1ε dynamics in the presence and absence of PRICKLE3. HEK T-REx 293 PRICKLE3 TetON cells (HA-PRICKLE3) were induced by ON treatment with doxycycline. Subsequently, cells were treated with 2.5 µM PF-670462 (PF-67) at various time points and analyzed by Western blot. β-ACTIN served as loading control. N = 5. B’) Densitometric quantification of Western blot (WB) band intensities. Results were normalized to the 0-hour time point. Statistical analysis was performed using linear regression, the corresponding *p*-values are reported, N = 5. C) Immunofluorescence imaging of membrane levels of VANGL1 and pVANGL2 in the presence of PRICKLE3. HEK T-REx 293 PRICKLE3 TetON cells were transfected with MYC-VANGL1 and CK1ε constructs. PRICKLE3 expression was induced overnight by treatment with doxycycline. D) Quantification of the membrane levels of VANGL and pVANGL. The average signal from the membrane protein was analyzed and normalized to the average signal from the cytoplasm. This normalization was done to account for variations in the transfection efficiency of VANGL1 and CK1ε, for more see Supp. Fig. 5C. Statistical analysis was performed using the SuperPlots of Data, the corresponding *p*-values are reported, N = 3 E) Protein *in vitro* kinase assay. Western blot showing phosphorylation assay performed using PRICKLE3 and CK1ε kinase. Reaction was carried out at two-time points (1 h, overnight (ON)) in the presence of ATP (2 mM) MgCl_2_ (10 mM). Subsequently, PRICKLE was detected by anti-PRICKLE3 and pan pSer/pThr antibody.

According to Feng et al., VANGL undergoes basal phosphorylation by CK1 in the endoplasmic reticulum prior to its transport to the plasma membrane, where it is subsequently hyperphosphorylated by CK1 (33). To elucidate the membrane levels of VANGL and phosphorylated VANGL in the presence of PRICKLE3 and CK1ε, we analysed the average signal from membrane-associated proteins and normalized it to the average signal from the cytoplasm (**Fig. 6C**, **Suppl. Fig. 5B,C**). This normalization was performed to account for variations in the transfection efficiency of VANGL1 and CK1ε. Our findings indicate that the presence of PRICKLE3 enhances the level of VANGL at the membrane while simultaneously reducing the pVANGL2/VANGL1 ratio (**Fig. 6D**, **Suppl. Fig. 5D**), suggesting that membrane-bound VANGL is less phosphorylated.

To further investigate the mechanism by which PRICKLE3 influences VANGL phosphorylation, we conducted an *in vitro* protein kinase assay. Notably, we observed that PRICKLE3 can undergo phosphorylation by CK1ε (**Fig. 6E,E’**). This suggests that a possible mechanism for the reduction in VANGL phosphorylation involves competition between VANGL and PRICKLE3 for CK1-mediated phosphorylation. This finding introduces a novel role for PRICKLE3 within the WNT/PCP pathway in a fine-tuning phosphorylation of the plasma membrane proteins crucial for non-canonical WNT signalling – RNF43 and VANGL1/2, which results in the VANGL1/2 stabilization at the cell periphery and suppression of the RNF43 enzymatic activity. Inhibition of RNF43 explains the potentiation of the WNT5A response and lack of the effect in the absence of the endogenous Wnt ligands (**Fig. 3B**).

In summary, we show that PRICKLE3 plays a role in both the WNT5A-dependent and WNT5A- independent, but CK1-dependent, aspects of WNT/PCP signaling (**Fig. 7**).

**Figure 7.**
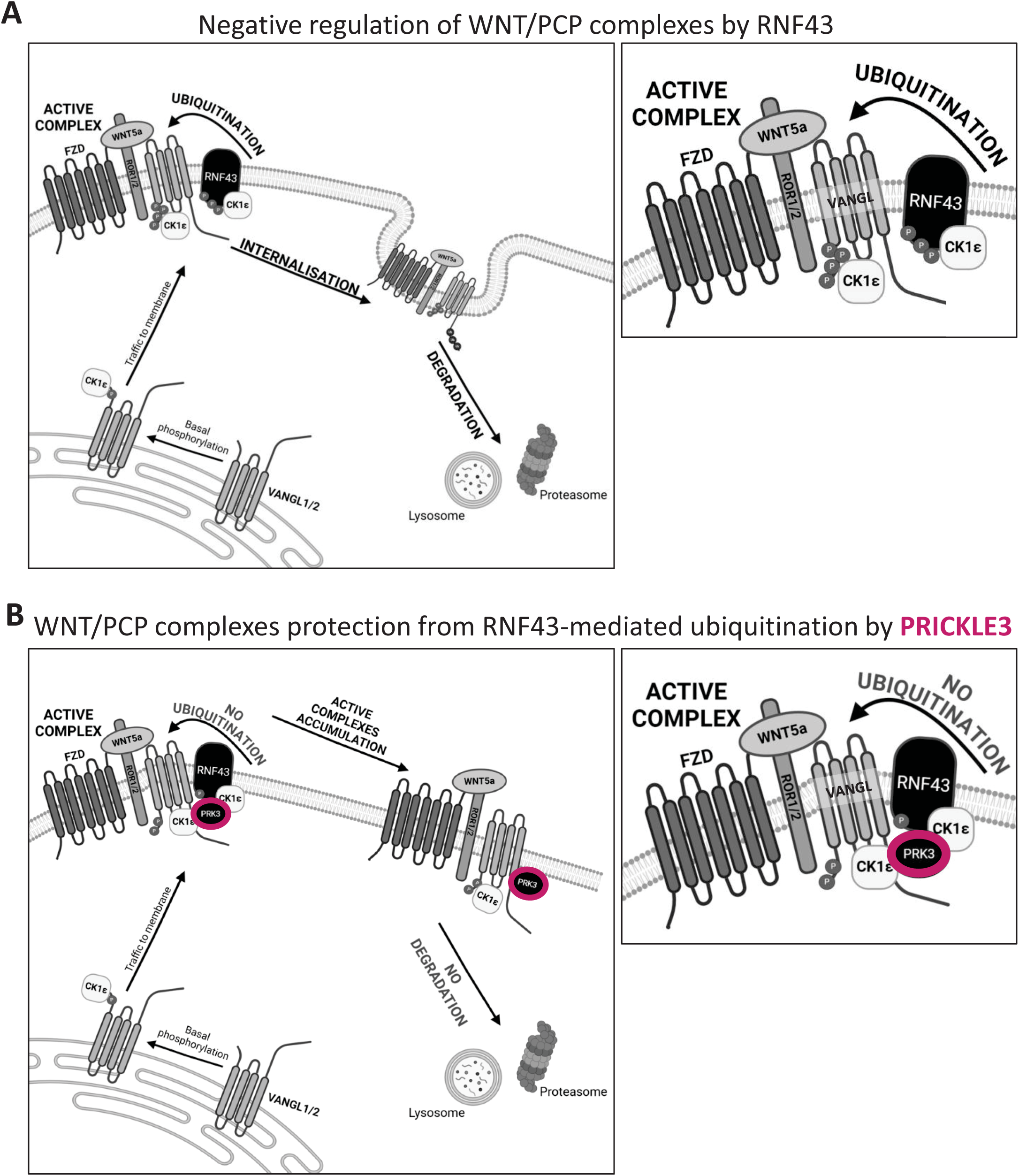
Summary of PRICKLE3 function, emphasizing its role and mechanisms of action in the stabilization of VANGL proteins. A) Negative regulation of WNT/PCP complexes by RNF43. Casein kinase 1ε plays a dual role in the regulation of WNT/PCP complexes. CK1ε phosphorylation is essential for the proper translocation of VANGL proteins to the plasma membrane. The activity of this kinase is also required for the function of the FZD and ROR, the WNT receptors and co-receptors. In addition, the enzymatic activity of the E3 ubiquitin ligase RNF43, a negative regulator of WNT signalling, is also promoted by CK1ε. As a result, WNT/PCP plasma membrane complexes are degraded. This relationship establishes a CK1ε/RNF43-mediated negative feedback loop that tightly controls the activation level of the PCP signaling pathway. Created withBioRender.com B) Protection from RNF43-mediated ubiquitination. PRICKLE3 disrupts this CK1ε/RNF43 feedback loop. It results in the stabilization of VANGL1/2 and ROR2 protein activation at the plasma membrane. Also, phosphorylation of membrane-bound VANGL1/2 is reduced; PRICKLE3 thus supports WNT/PCP signaling. Created with BioRender.com

## DISCUSSION

This report is the first to utilize the miniTurboID assay for a direct comparison of PRICKLE1-3 isoforms at the interactome level. Through this approach, we confirmed previously known interactors of PRICKLE proteins and uncovered potential novel interactions. For instance, Misshapen-like kinase 1 and CK1 kinases, which were previously described as interactors of PRICKLE1 (55–57), were identified for PRICKLE1-3. Interestingly, all our baits significantly neighbour with CK1δ/ε but also with CK1α, which was previously not reported. Based on our datasets, PRICKLE3 can interact with SRCIN1 (also known as SNAP-25-Interacting Protein) which was shown to destabilize β-catenin and affect the composition of the immune tumor microenvironment in breast cancer (58) and be involved in the Src and STAT3 signaling in neuroblastoma (59). *Xenopus laevis* homolog of the other protein found here – *epbl41l4a* is expressed in the rotating somites and the forming neural tube, suggesting involvement in the WNT/PCP pathway (60). These are just examples of the significance of our work and potential future directions set by our study.

Our findings indicate that interactors of PRICKLE1-3 are quite uniformly localized within cells – primarily at the periphery, centrosomes, endomembrane system, and mitochondrial matrix. However, many of the identified interactors were not mapped in the Human Cell Map resources, as the authors used the less robust BirA*-based BioID protocol. Notably, the isoform-specific proximal interactome of PRICKLE1 is enriched in centrosome, while PRICKLE2 in mitochondrial components. Interestingly, to date, only PRICKLE3 has been linked to this organelle, as it has been associated with mitochondrial dysfunction and ATPase biogenesis (61,62). Additional investigation is needed to understand how PRICKLE2 signals to mitochondria.

Further, we decided to focus more on the isoform-specific interactions. Already at the early stage of the work with generated data, PRICKLE3 seemed to be different from PRICKLE1/2. Next, analysis revealed that the proximal microenvironment of the PRICKLE3 is enriched in the proteins forming GO term dedicated to the WNT/PCP pathway. The following analysis revealed that PRICKLE3, fused with miniTurboID enzyme, marked the most significant components of the non-canonical WNT signalling, including VANGL1/2, CELSR1/2, KIF26B, ROR1/2, and FZD3 (16,18,27). This directly suggested involvement in the signalling events occurring at the plasma membrane, which was subsequently experimentally validated. In line with that, we identified functional distinctions in non-canonical WNT signaling regulation between PRICKLE1 and PRICKLE3. Notably, PRICKLE1 showed no activity in our model, while PRICKLE3 acted as a positive regulator of the pathway as it stabilized VANGL by interfering with RNF43 and CK1ε enzymatic functions and supported ROR2 shift in a WNT5A-dependent manner.

The role of CK1 kinases in protein degradation is well-established, with CK1α marking β- catenin and CK1δ targeting MDM2 for proteasomal degradation (63–65). Building on previous findings by Feng et al., which demonstrated that Vangl proteins are predominantly degraded via the proteasomal pathway (33), we concentrated our investigation on this specific degradation mechanism. In contrast to Feng et al., who focused on the cytoplasmic pool of Vangl and its endoplasmic reticulum-associated degradation, our research examines the degradation of Vangl at the plasma membrane. Here, we explore CK1-mediated regulation of protein turnover through previously described mechanisms, where the enzymatic activity of the E3 ubiquitin ligase RNF43 is promoted by CK1-mediated phosphorylation. Using a variety of assays, we expanded the current understanding of the mechanism by which RNF43 negatively regulates WNT5A/PCP receptor complexes (31). Specifically, we demonstrate that PRICKLE3 can interfere with RNF43 activation by CK1, suggesting that PRICKLE3 competes with RNF43 for access to CK1 enzymatic activity. This finding aligns with previous observations showing that Prickle2, in complex with Vangl2, undergoes polyubiquitination (38). We also see the ubiquitination of the PRICKLE3 in presence of VANGL2 and RNF43. Thus, it is likely that PRICKLE3 presence limits VANGL ubiquitination by RNF43 functional saturation.

Interestingly, our findings reveal that excessive CK1 activity can inhibit the non-canonical WNT pathway. Specifically, we observed that the VANGL protein remains more stable at the plasma membrane when CK1-mediated phosphorylation is moderate. This study both supports and extends existing knowledge on CK1-mediated trafficking of VANGL to the plasma membrane. Notably, our findings do not contradict previous research indicating that the non-phosphorylatable VANGL mutant is unstable (33,34). Here, we propose that the interaction between PRICKLE3 and VANGL occurs specifically at the plasma membrane. Prior studies have suggested that excessive phosphorylation within the N-terminus of Vangl can lead to its destabilization at the plasma membrane in *Drosophila melanogaster* cells, and that Prickle plays a protective role by preventing Vangl phosphorylation and subsequent degradation in this model organism (44,66). Our research confirms these findings in a human cell line. Additionally, our *in vitro* phosphorylation assay directly demonstrates that PRICKLE3 can be phosphorylated by CK1ε, providing further support for our data.

In the fruit fly Prickle, phosphorylated by Nemo kinase, is marked to be degraded in proteasome by the E3 ubiquitin ligase complex Cullin1(Cul1)/SkpA/Supernumerary limbs (Slimb) (40,67). This suggests the existence of the evolutionary conserved mechanism of PCP regulation by Prickle protein level control via phosphorylation, ubiquitination and proteasomal degradation. Interestingly, we found Ubiquitin Specific Peptidase 31 (USP31) that upregulated in our datasets. This can potentially act as PRICKLE3 deubiquitinase.

Interestingly, we did not observe negative regulation of the Dishevelled isoforms, which contrasts with previous studies (41,42,45). Specifically, we found no antagonistic effect on the levels of DVL2/DVL3 proteins or their phosphorylation states. These discrepancies may stem from various factors, such as overexpression artifacts (68), differences in model organisms (e.g., *Drosophila melanogaster*, which lacks an RNF43 homolog (69), or variations in antibody specificity. Specifically, the WB band produced by the anti-DVL3 antibody (sc-8027, Santa Cruz Biotechnology) can differ, depending on the DVL3 phosphorylation level (70). In our study, we monitored endogenous DVL2/3 using specific antibodies in clonal PRICKLE1/3 Tet ON cell lines. This approach helped mitigate transfection-related stress and variability associated with transfection efficiency. Therefore, our findings provide a clearer picture of Dishevelled regulation under more physiological conditions.

Finally, the dataset generated in this study has been rigorously validated, offering detailed insights into the cellular microenvironment of PRICKLE isoforms. We believe this dataset serves as a valuable resource for advancing the understanding of PRICKLE isoform functions in human cells, particularly in the context of cellular signaling and structural dynamics. Notably, we have established a functional connection between PRICKLE3 and RNF43, an E3 ubiquitin ligase known for its crucial regulatory roles in development and tumorigenesis (71,72). RNF43 has been widely implicated in cancers of the gastrointestinal tract, skin, ovaries, and other tissues due to its role in modulating WNT signaling, a pathway essential for both normal cellular growth and oncogenesis. Dysregulation of RNF43 disrupts this pathway, contributing to tumor progression and metastasis (72,73). Our findings underscore the potential of PRICKLE3 to influence RNF43 activity, providing a new avenue for exploring PRICKLE3’s impact on WNT-related signaling pathways in both developmental and disease contexts.

## ACKNOWLEDGEMENTS

We would like to thank the members of Harnos lab and Štěpán Čada for a fruitful discussion, and Lenka Doubková and Dr. Eva Slabáková for excellent administrative support. We extend our sincere gratitude to Dr. Konstantinos Tripsianes and Dr. Sara Bologna for providing the purified recombinant CK1ε protein.

## AUTHOR CONTRIBUTIONS

K.A.R. had the main ideas, performed key experiments, analyzed the data, drafted the manuscript, performed a literature search, and made the figures.

T.W.R. performed experiments and greatly helped with manuscript preparation and shaping of the whole project.

P.K. and P.P. performed additional experiments with cell lines.

K.G. performed all bioinformatic analyses.

T.B. was involved in the preparation of cell lines.

K.H. performed the mass spectrometry experiments and prepared KNIME workflow

Z.Z. supervised the mass spectrometry experiments.

J.H. critically revised the manuscript and figures, secured the funding, and supervised the overall work.

## MATERIALS AND METHODS

### Cell culture

All cell lines utilized in this study (**Suppl. Table 10**) were cultured in Dulbecco’s Modified Eagle Medium (DMEM, 41966-029, Gibco, Life Technologies), supplemented with 10% fetal bovine serum (FBS, 10270-106, Gibco, Life Technologies) and 1% penicillin-streptomycin (XC-A4122/100, Biosera). The cells were maintained at 37°C in a 5% CO_2_ humidified atmosphere. Routine tests for mycoplasma contamination were regularly conducted to ensure the absence of contamination in the cell cultures.

### Cells treatments

Cells with a doxycycline-inducible system were treated with 0.1- 1μg/ml of doxycycline (HY-N0565B, MedChem Express) overnight to induce protein expression.

To inhibit endogenous WNT ligands, cells were treated with 1 μM LGK-974 (1241454, PeproTech) overnight. Following this, non-canonical WNT signaling was activated by incubating the cells with recombinant human WNT5A (645-WN, R&D Systems) at a concentration of 100 ng/mL for 3 hours. Similarly, canonical WNT signaling was activated by incubating the cells with 100 ng/mL of recombinant human WNT3A (5036-WN, R&D Systems) for 3 hours.

To assess protein degradation kinetics, cells were pretreated with doxycycline overnight. The following day, the cells were treated with 20μM cycloheximide (Sigma-Aldrich, C7698) and harvested at various time points for analysis.

To inhibit CK1ε activity, cells were treated with 2.5 μM PF670462 (sc-204180, Santa Cruz,) for the indicated time.

### Preparation of stable cell lines

HEK239 T-REx PRICKLE3 and PRICKLE1 TetON cell lines were generated by transfecting HEK239 T-REx cells (R71007, Invitrogen) with the pcDNA4TO-zeo plasmid encoding N-terminally HA-tagged PRICKLE isoforms. Similarly, HEK293 T-REx miniTurboID TetON cell lines were established by transfecting HEK293 T-REx cells with pcDNA4/TO-zeo plasmids encoding human PRICKLE1-3 proteins, tagged at the N-terminus with miniTurboID biotin ligase and a V5 tag, or with a control plasmid containing only the miniTurboID enzyme and a V5 tag (details in **Suppl. Table 11**). Twenty-four hours post-transfection, cells were selected using 100 µg/mL of Zeocin (ant-zn-1p, Invivogen). Subsequently, single clones were isolated, propagated, and validated through Western blot analysis and immunofluorescence.

### Plasmids/cloning

The generation of all plasmids involved the amplification of sequences using Q5® High-Fidelity DNA Polymerase (M0491S, New England Biolabs), followed by cloning utilizing the In-Fusion cloning method (639690, Takara Bio). All plasmids utilized in this study are listed in **Suppl. Table 11**.

To generate the TetON-inducible pcDNA4-HA-PRICKLE3 plasmid, the cDNA encoding PRICKLE3 was amplified from the pGFP_LMO6 (PRICKLE3) plasmid. For the construction of the TetON-inducible pcDNA4-HA-PRICKLE1 plasmid, the cDNA encoding PRICKLE1 was amplified from the pCMV6-XL5- Prickle1 plasmid. The amplified sequences were subsequently cloned into the pcDNA4-TO-RNF43- 2xHA-2xFLAG plasmid, which had been linearized using HindIII (ER0501, Thermo Fisher Scientific) and XbaI (ER0681, Thermo Fisher Scientific).

For the generation of the pcDNA4-CTRL-miniTurboBioID plasmid, the cDNA encoding V5-miniTurboID was amplified from the pCW57-EPHA2-V5-miniTurboID plasmid. Similarly, to generate the pcDNA4- PRICKLE1-3 miniTurboBioID plasmids, the cDNA encoding V5-miniTurboID was amplified from the pCW57-EPHA2-V5-miniTurboID plasmid, and the PRICKLE1-3 sequences were amplified from the pCMV6-XL5-Prickle1, pGateway 3XFlag Prickle2, and pGFP_LMO6 (PRICKLE3) plasmids, respectively. The resulting sequences were cloned into the pcDNA4-TO-RNF43-2xHA-2xFLAG plasmid which had been linearized using HindIII and XbaI.

The pcDNA3-RNF43-FLAG and pcDNA3-RNF43 Ring mut-FLAG plasmids were generated by PCR skipping the HA tag from the pcDNA4-TO-RNF43-2xHA-2xFLAG and pcDNA4-TO-RNF43Mut1-2xHA- 2xFLAG plasmids, respectively. The sequences were then cloned into pcDNA3, which had been linearized using HindIII and EcoRV (ER0301, Thermo Fisher Scientific).

### Transfection

Cells were transfected using the jetOPTIMUS transfection reagent (117-07, Polypus) at a 1:1 ratio with plasmid DNA, or alternatively, transfections were performed using 1 μg/ml polyethyleneimine (PEI) (23966-100, Polysciences) at pH 7.4, with plasmid DNA in a 4:1 ratio. The culture media were replaced 6 hours post-transfection. The amount of plasmid DNA used for transfection was as follows: 300 ng per well of a 24-well plate for antibody verification, 2.5 µg for a 4 cm culture dish for ubiquitination assays, 10 µg for a 10 cm dish for co-immunoprecipitation or stable cell line preparation.

### Western-blot analysis

Cells were lysed directly in a buffer containing 100 mM Tris/HCl (pH 6.8), 20% glycerol, 1% SDS, 0.01% bromophenol blue, and 1% 2-mercaptoethanol. Protein separation by SDS-PAGE was performed according to the manufacturer’s protocol with minor modifications (Bio-Rad). Briefly, proteins were resolved on an SDS-PAGE gel at 110 V and subsequently transferred to an Immobilon-P PVDF membrane (Millipore, IPVH00010) for 1 hour at 100 V. Membranes were then blocked in a 5% non-fat dry milk solution prepared in TBS-T for 30 minutes and incubated overnight at 4 °C with primary antibodies. Following incubation, membranes were washed in TBS-T and treated with HRP-conjugated secondary antibodies. Protein detection and visualization were carried out using the Fusion SL imaging system (Vilber) with the Immobilon Western Chemiluminescent HRP Substrate (Merck, WBKLS0500). A complete list of antibodies used in this study is provided in **Suppl. Table 12**.

### Immunoprecipitation

Cells were transfected with the appropriate plasmid DNA and/or induced with doxycycline overnight and cultured for 24 hours. Subsequently, the cells were washed once with PBS and lysed for 5 minutes in a cold buffer consisting of 50 mM Tris (pH 7.6), 200 mM NaCl, 1 mM EDTA, 0.5% NP40, fresh 0.1 mM DTT (E3876, Sigma), a protease inhibitor cocktail (04693159001, Roche), and a phosphatase inhibitor (No. 524625, Merck). The samples were then sonicated. Cell debris was removed by centrifugation at 16,000 g for 15 minutes at 4°C, and 10% of the total cell lysate was preserved as a control for Western blot analysis. Lysates were incubated with 1 μg of antibody for 16 hours at 4°C on a head-over-tail rotator. Subsequently, 20 μL of protein G-Sepharose beads (17-0618- 05; GE Healthcare), equilibrated in complete lysis buffer, were added to each sample and gently rotated for 6 hours at 4°C. After this incubation, the samples were washed six times with lysis buffer and then resuspended in 100 μL of Western blot sample buffer. The immunoprecipitation experiments were analyzed using Western blotting.

### His-ubiquitin pulldown assay

The His-ubiquitin pulldown assay was performed as described by Radaszkiewicz et al. (31).

Briefly, HEK239 T-REx PRICKLE3 TetON cells were transfected with plasmids encoding polyhistidine-tagged ubiquitin, EGFP-VANGL2, and FLAG-tagged wild-type or enzymatically inactive Mut1 RNF43 constructs. Six hours post-transfection, the culture media were replaced, and PRICKLE3 expression was induced with doxycycline. Twenty hours after transfection, the cells were treated with 5 μM MG- 132 (HY-13259, MedChem Express) for 5 hours. Subsequently, the cells were lysed in a buffer containing 6 M guanidine hydrochloride (G3272, Sigma) 0.1 M NaxHxPO4 pH 8.0, and 10 mM imidazole (I5513, Sigma), then sonicated and boiled. Samples were centrifuged at 16,000 g for 10 minutes at room temperature to remove insoluble fractions. Ten percent of the cellular lysate was retained as a transfection control, which was subjected to ethanol precipitation and subsequently resuspended in the Western blot sample buffer. To perform the pull-down of tagged proteins, 10 µL of equilibrated His Mag Sepharose Ni beads (GE28-9799-17, GE Healthcare) were added to each sample and incubated on a roller for 6 hours at room temperature. The beads were subsequently washed three times with a buffer composed of 8 M urea (U5378, Sigma), 0.1 M NaH2PO4 (pH 6.3), 0.01 M Tris, and 15 mM imidazole. Following the washes, the beads were resuspended in 100 μL of Western blot sample buffer, boiled for 5 minutes, and loaded onto an SDS-PAGE gel for further analysis.

### *In vitro* kinase assay

An *in vitro* kinase assay was performed using full-length human CK1ε (kindly provided by Dr. Konstantinos Tripsianes and Sarah Bologna) (74) and recombinant PRICKLE3 (Antibodies Online, ABIN1316198) as the substrate in a buffer containing 25 mM Tris-HCl (pH 7.5), 250 mM NaCl, 1 mM DTT, 10 mM MgCl2, 1 mM EDTA, and 2 mM ATP. The molar ratio was set to 1:10, with 0.21 μM substrate and 2.1 μM kinase in a 25 μl reaction mix. A sample collected immediately after initiation of the reaction (upon MgCl2 addition) served as the control. The reaction was then carried out for 1 hour and overnight. Reactions were stopped by adding 2× Laemmli buffer and boiling at 95 °C for 1 minute. Samples were subsequently analyzed by WB.

### Immunofluorescence and confocal microscopy

Cells were plated on glass coverslips that had been coated for 30 minutes at 37°C with Cultrex (3536- 001-02, Biotechne, R&D Systems) diluted 1:500 in PBS. The cells were fixed with 4% paraformaldehyde (PFA) (1.00196, Sigma-Aldrich), in PBS, followed by permeabilization with 0.1% Triton X-100 in PBS and blocking with a 1% solution of bovine serum albumin (BSA) in PBS. The samples were then incubated overnight at 4°C with primary antibodies diluted in 1% BSA in PBS. The following day, the cells were incubated with the appropriate Alexa Fluor™ secondary antibodies (Invitrogen) for 1 hour at room temperature, also diluted in 1% BSA in PBS. To visualize biotinylation reaction products, streptavidin-Alexa Fluor™ conjugate was utilized. Nuclei were counterstained with 1 μg/mL DAPI (62248, Invitrogen). Finally, the samples were mounted using DAKO medium (S3023, DAKO). Images were acquired using a Leica TCS SP8 confocal laser scanning microscopy platform (Leica).

### Image analysis of the membrane levels of protein

Image analysis was conducted using ImageJ software. To evaluate the membrane levels of proteins, we analyzed the average signal from membrane-associated proteins and normalized it to the average signal from the cytoplasm. Briefly, regions of interest (ROIs) were defined to encompass both the membrane and cytoplasmic areas (see **Suppl. Fig. 5C**). We measured the gray value intensity and quantified the mean intensity values for both membrane and cytoplasm. The ratio of these values represented the membrane levels of the protein. Three independent experiments were performed, with grey value intensity measured in at least 40 cells for each condition in every experiment. Statistical analysis of the results and graph preparation were conducted using the SuperPlots of Data tool (75) available at https://huygens.science.uva.nl/SuperPlotsOfData/.

### miniTurboID samples preparation

A miniTurboID assay was performed according to the published protocol (76) with minor modifications (details to follow). HEK293 T-REx miniTurboID TetON cell lines were cultured in 15 cm dishes. Once the cells reached confluency, they were treated with doxycycline (1 μg/mL) overnight to induce protein expression. Following doxycycline induction, the cells were stimulated with biotin (50 μM; Santa Cruz Biotechnology) for 6 hours. The cells were subsequently washed twice with PBS and lysed on ice for 15 minutes in a cold lysis buffer consisting of 50 mM Tris (pH 7.6), 500 mM NaCl, 0.4% SDS, and freshly added: 2% Triton X-100, 1 mM DTT (E3876, Sigma), and a protease inhibitor cocktail (04693159001, Roche). The lysates were gently sonicated and incubated on ice for 5 minutes. Cellular debris was removed by centrifugation at 16,000 g for 15 minutes at 4°C. The supernatant was diluted two-fold with 50 mM Tris-HCl (pH 7.4). Ten percent of the total lysate was set aside for subsequent Western blot analysis. The remaining lysates were incubated with Streptavidin Sepharose High-Performance beads (17-5113-01, GE Healthcare) at 4°C with gentle agitation overnight to capture biotinylated proteins, using 40 μL of bead slurry per sample. After incubation, the samples were washed four times with a wash buffer containing 50 mM Tris (pH 7.6), 250 mM NaCl, 0.2% SDS, and 1% Triton X-100 (freshly added), followed by two additional washes with 50 mM Tris (pH 7.6). Finally, the beads were stored at −80°C for further analysis.

### Mass spectrometry data analysis (miniTurboID)

Proteins were extracted from the beads using 2% SDS solution for 20 mins in a thermomixer (Eppendorf ThermoMixer C, 20 min, 50°C, 750 rpm). After that, the beads suspension was centrifuged (2 min, 1,000 x *g*) and the supernatant was transferred to clean test tube. Dithiothreitol (0.5M solution in water) was added to the protein solution (final concentration 0.1M) and proteins were reduced for 15min at 95°C. Reduced proteins solution was used for filter-aided sample preparation (FASP) as described elsewhere (77) using 0.75 μg of trypsin (sequencing grade; Promega). The resulting peptides were extracted into LC-MS vials by 2.5% formic acid (FA) in 50% acetonitrile (ACN) and 100% ACN with the addition of polyethylene glycol (final concentration 0.001%) (78) and concentrated in a SpeedVac concentrator (Thermo Fisher Scientific).

LC-MS/MS analyses of all peptides were done using UltiMate 3000 RSLCnano system connected to Orbitrap Exploris 480 spectrometer (Thermo Fisher Scientific). Before LC separation, tryptic digests were online concentrated and desalted using a trapping column (Acclaim PepMap 100 C18, dimensions 300 μm ID, 5 mm long, 5 μm particles, Thermo Fisher Scientific). The trap column was then washed with 0.1% FA and the peptides were eluted in backflush mode from the trapping column onto an analytical column tempered to 50 °C (Aurora C18, 3rd generation, 25cm long, 75μm ID, 1.7μm particles, P/N AUR3-25075C18-TSI, Ion Opticks) by 60 min gradient program (flow rate 150 nl.min^-1^, 3-37% of mobile phase B; mobile phase A: 0.1% FA in water; mobile phase B: 0.1% FA in 80% ACN) followed by 7 min long system wash using 80% of mobile phase B. Equilibration of the trapping column and the analytical column was done before sample injection to sample loop. Spray voltage and sweep gas were set to 1.6kV and 1, respectively.

Data were acquired in a data-independent acquisition mode (DIA). The survey scan covered m/z range of 350-1400 at a resolution of 60,000 (at m/z 200) and a maximum injection time of 55 ms (normalized AGC target 300%). HCD MS/MS (27% relative fragmentation energy) were acquired in the range of m/z 200-2000 at 30,000 resolution (maximum injection time 55 ms, normalized AGC target 1000%). Overlapping windows scheme in the precursor m/z range from 400 to 800 were used as isolation window placements – see **Suppl. Table 13** for more details. Raw data were converted to mzML format using msconvert (version 3.0.21193-ccb3e0136) using peakPicking (vendor msLevel=1-) and demultiplex (optimization=overlap_only massError=10ppm) filters applied.

DIA data in mzML format were processed in DIA-NN (79) (version 1.8.1) in library free mode against a modified cRAP database (based on http://www.thegpm.org/crap/; 111 sequences in total) and UniProtKB protein database for *Homo sapiens* (https://ftp.uniprot.org/pub/databases/uniprot/current_release/knowledgebase/reference_proteomes/Eukaryota/UP000005640/UP000005640_9606.fasta.gz version from 2023-11-08, number of protein sequences: 20,596). No optional, but carbamidomethylation as fixed modification and trypsin/P enzyme with 1 allowed missed cleavages and peptide length 7-30 were set during the library preparation. False discovery rate (FDR) control was set to 1% FDR. MS1 and MS2 accuracies as well as scan window parameters were set (8, 21, 9, respectively) based on the initial test searches (median value from all samples ascertained parameter values). MBR was switched on.

The MaxLFQ intensities, reported in the output of DIA-NN, report.tsv were further processed using the software container environment (https://github.com/OmicsWorkflows), version 4.7.7a. The full processing workflow is available on the WorkflowHub under the identifier 1202 (https://workflowhub.eu/workflows/1202) with DOI 10.48546/WORKFLOWHUB.WORKFLOW.1202.1. Briefly, it covered a) removal of low-quality precursors and contaminant protein groups, b) protein group intensities log2 transformation and LoessF normalization, c) imputation of missing values by random draws from a manually defined left-shifted Gaussian distribution originated from the actual data with shift of 1.8 and scale 0.3 in a column-wise manner, d) differential expression analysis using LIMMA statistical test calculated for protein groups.

Follow-up analyses were carried out in R, v. 4.4.0 and are available in the GitHub repository: https://github.com/HarnosLab/2024_Radaszkiewicz, with DOI 10.5281/zenodo.14142009. Data visualizations were done using ComplexHeatmap, UpsetR, ggplot2 R packages, gene ontology was done using gProfiler2 (version *e111_eg58_p18_f463989d*) and Metascape tools. Saint probability score for bait-prey interactions was computed using the REPRINT resource (26), v. 2.0, with the following settings: experiment type was set to Proximity Dependent Biotinylation; File Type as tab-separated matrix; only user controls were selected. To compute the probabilistic SAINT score, SAINTexpress was used. DotPlots were created using the ProHits-viz suite (80).

The mass spectrometry proteomics data have been deposited to the ProteomeXchange Consortium via the PRIDE partner repository with the dataset identifier PXD057854. <u>Reviewer access details</u>:

Log in to the PRIDE website using the following details:

Project accession: PXD057854
Token: 9XuZULOPc9UY

Alternatively, reviewer can access the dataset by logging in to the PRIDE website using the following account details:

Username:
Password: LLJblukqqWsQ.

### Software and statistics

Statistical significance was determined using two-tailed unpaired Student’s t-tests or linear regression analysis. All statistical details including the number of biological or technical replicates can be found in each figure legend. Statistical analysis and data visualization were performed in GraphPad Prism 8.0 software. Graphs are presented with error bars as ± SD if not stated differently in the figure legends. Figures 1a, 2a, 2b, and 7 were created with BioRender.com, the online app used to create, edit, and collaborate on scientific diagrams, and illustrations.

## ABBREVIATIONS

BSA: Bovine serum albumin
CELSR: Cadherin EGF LAG seven-pass G-type receptor
CHX: Cycloheximide
CK1: Casein kinase I
DOX: Doxycycline
DVL: Dishevelled
FZD: Frizzled
GO: Gene Ontology
HCM: Human Cell Map
KO: knockout
LRP6: Co-receptor low-density lipoprotein receptor-related protein 6
MS: Mass spectrometry
PCA: Principal component analysis
POI: Protein of interest
RNF43: E3 ubiquitin-protein ligase Ring finger protein 43
ROI: Regions of interest
ROR: Receptor tyrosine Kinase-like Orphan Receptor
SDS-PAGE: Sodium dodecyl sulfate–polyacrylamide gel electrophoresis
USP31: Ubiquitin Specific Peptidase 31
VANGL: Van Gogh-like
VBM: Vangl binding motif
WB: Western Blot
WNT/PCP: WNT/planar cell polarity
WT: Wild type

## Supplementary Materials

This Supplementary Material file contains 5 Figures and 13 Tables.

**Supplementary Figure 1.**
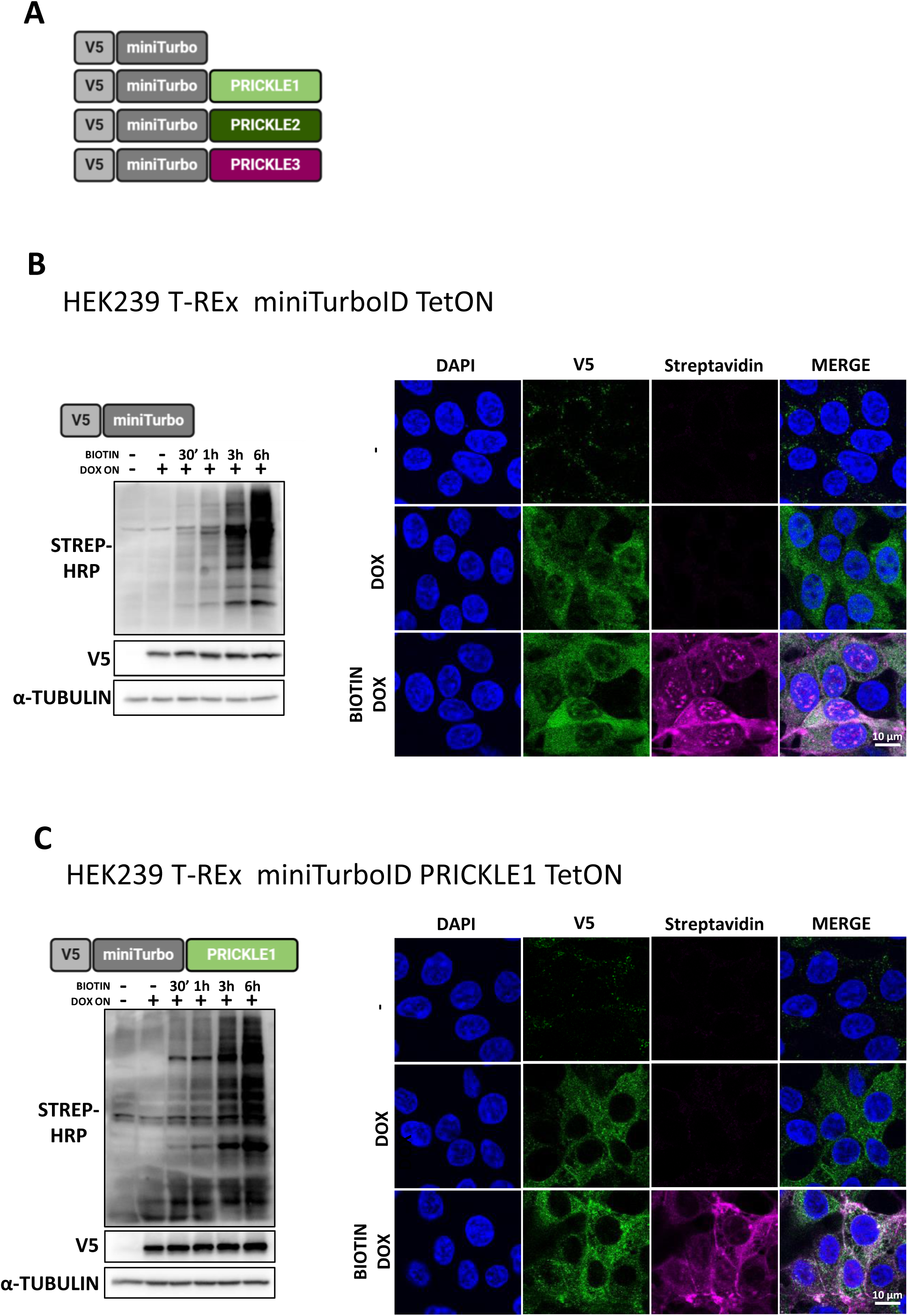
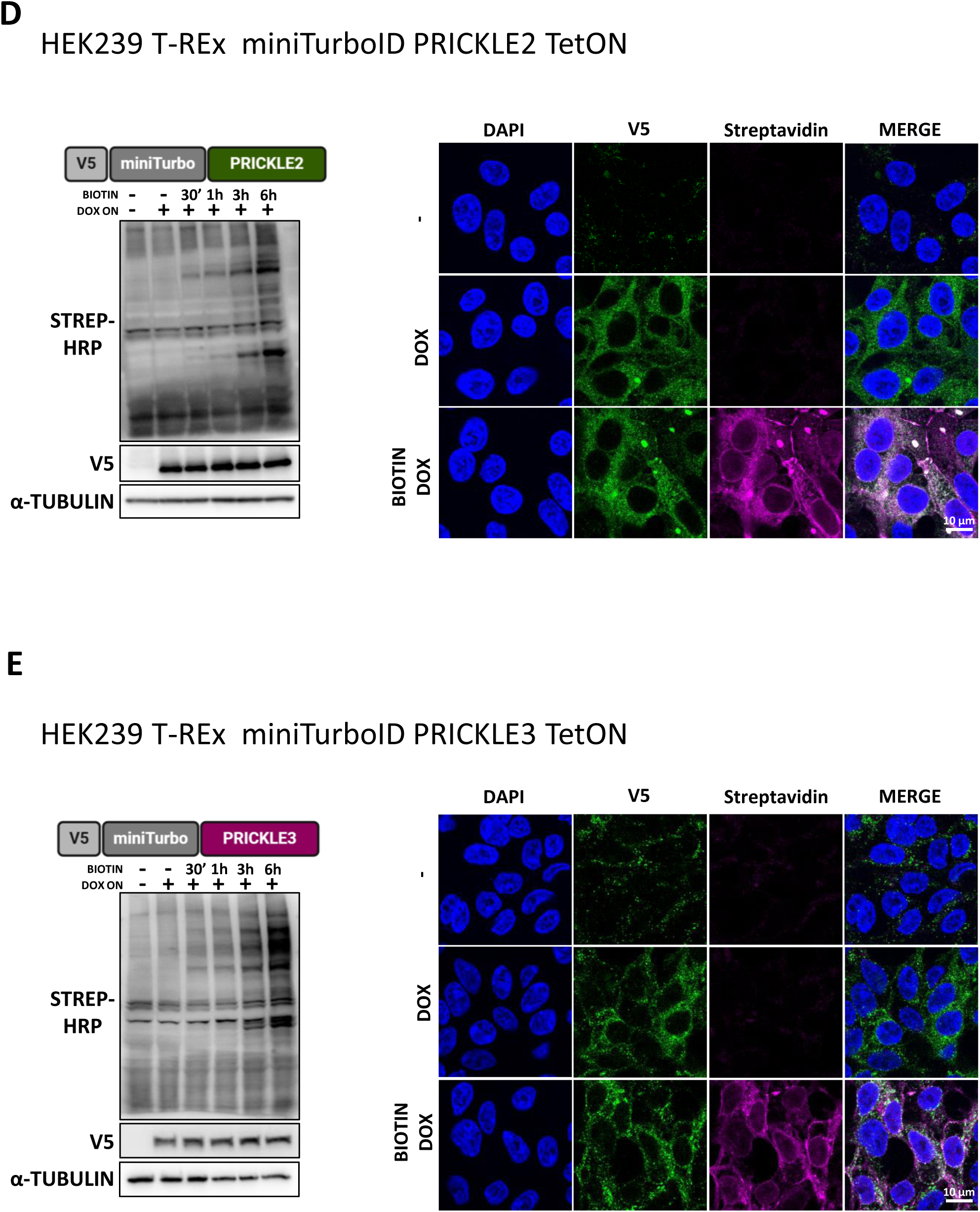
**A)** Schematic representation of the constructs. Human PRICKLE1-3 proteins were tagged at the N-terminus with the miniTurboID and V5 tags. A construct containing only the miniTurboID and V5 tag was used as the control. **B-E)** Validation of HEK T-REx 293 miniTurboID cell lines (B – control; C – PRICKLE1; D – PRICKLE2; E – PRICKLE3). To confirm expression induction and specificity of the biotinylation reaction, cell lines were treated with doxycycline overnight (dox ON). Biotin was subsequently added at various time points to initiate the enzymatic reaction. Samples were then analyzed by Western blot (WB) to assess biotinylation levels and baits expression (V5 tag). The α-TUBULIN protein level was used as loading control for WB. Additionally, cell lines were tested by immunofluorescence to analyse the location of baits upon treatment with biotin and doxycycline. Streptavidin was used for biotinylation reaction product visualization, V5 antibody to visualise construct expression, and nuclei were stained with DAPI.

**Supplementary Figure 2.**
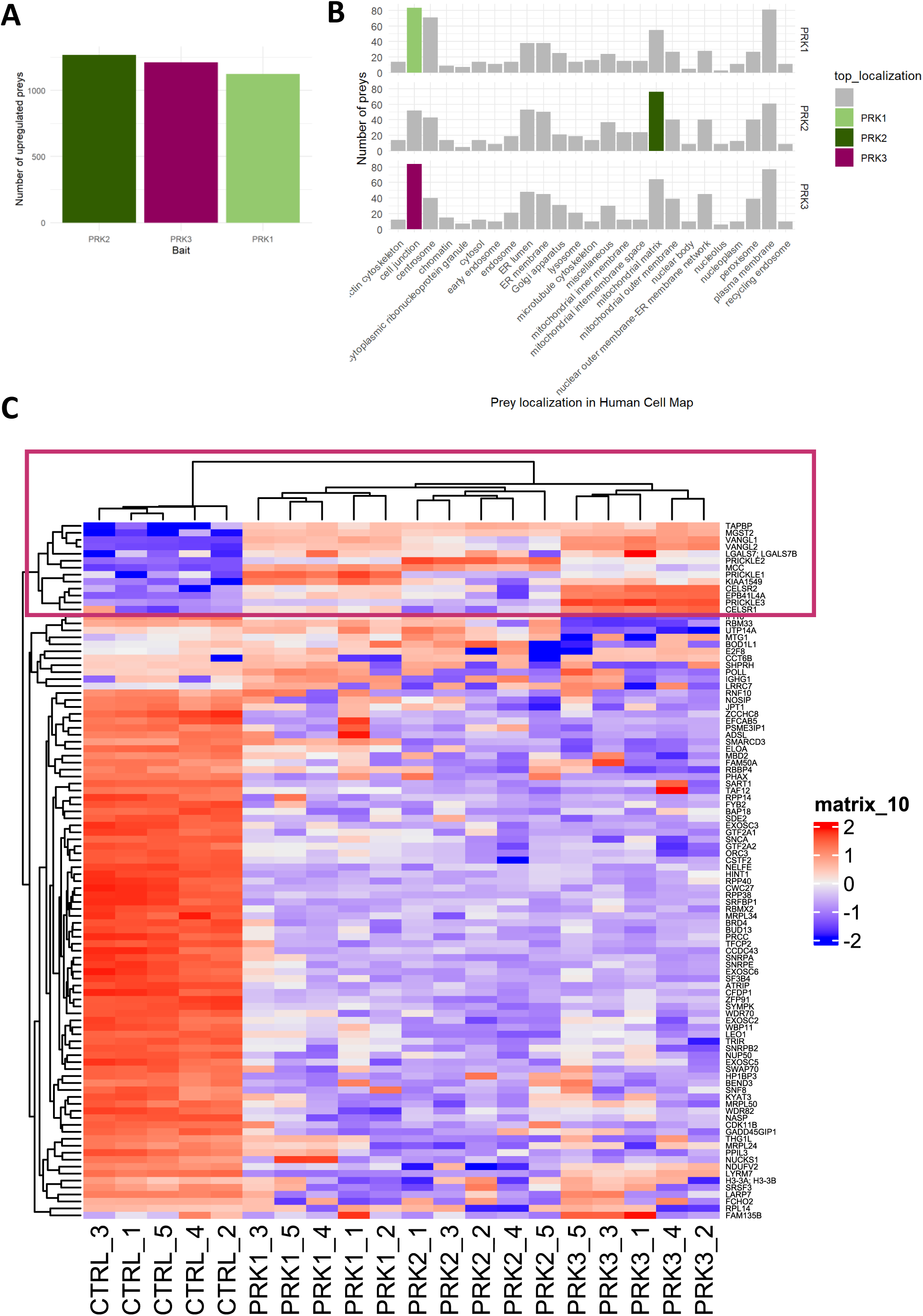
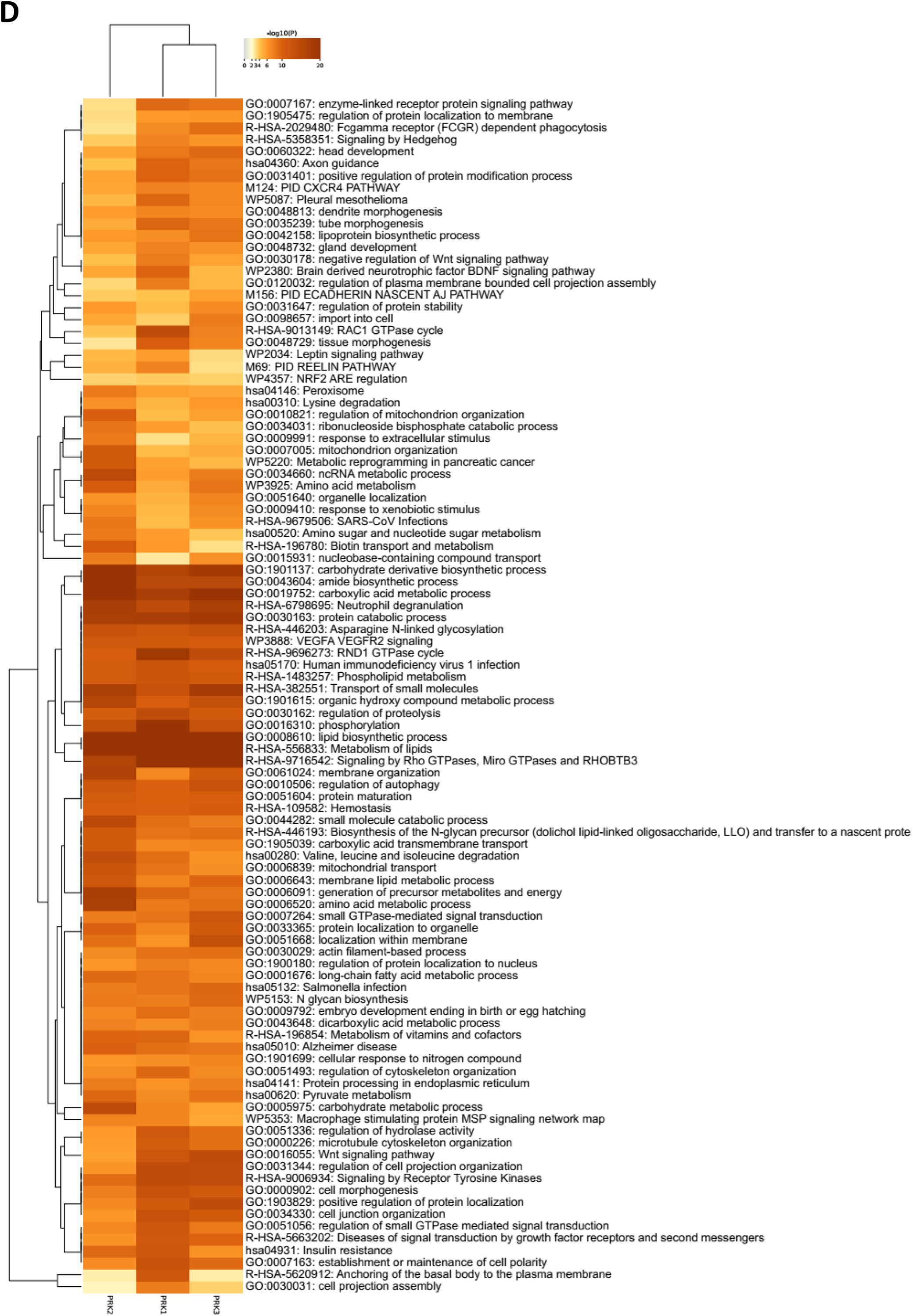
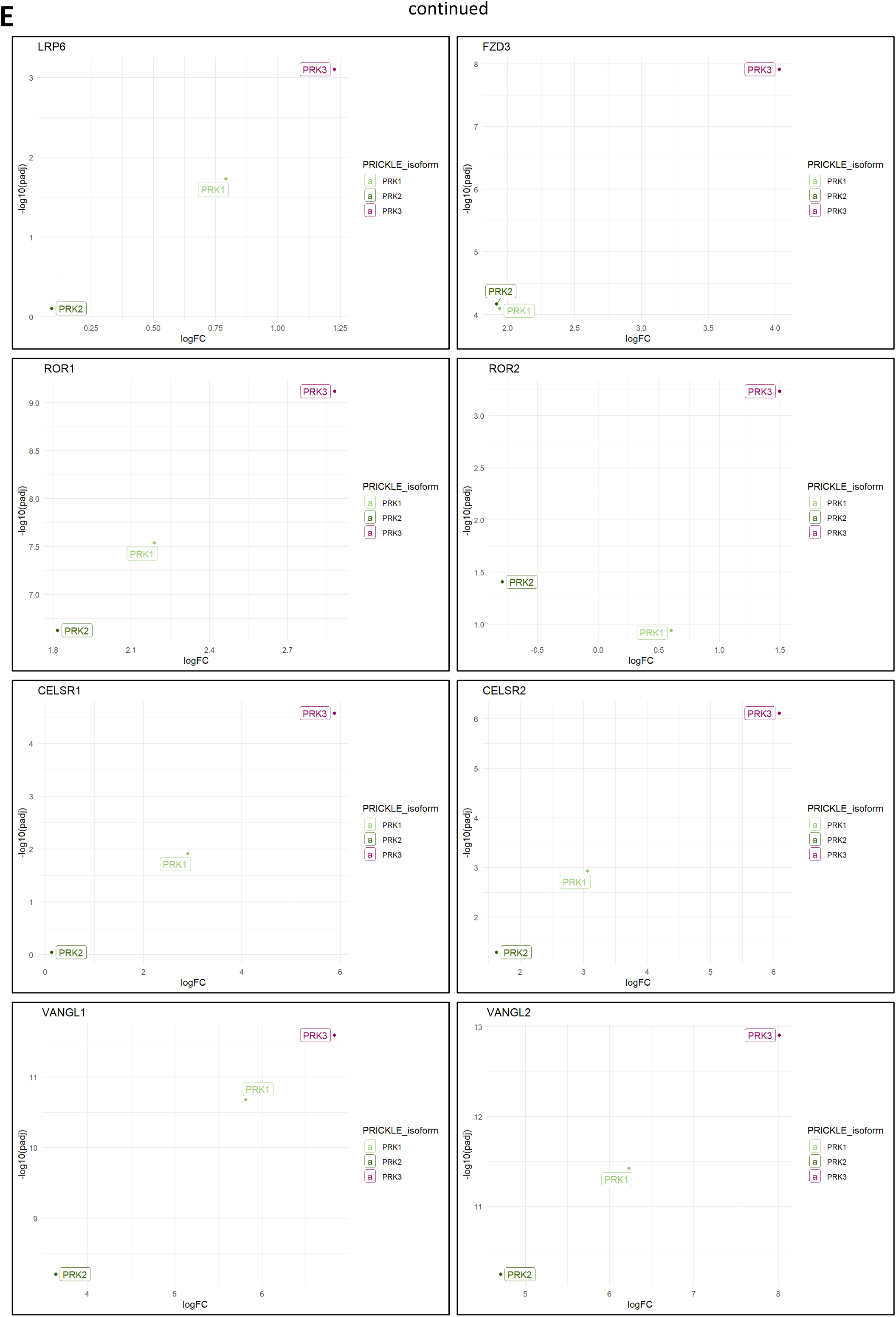
**A)** The number of upregulated prey proteins for each bait, which was determined based on a log2 fold change greater than 1 and an adjusted p-value less than 0.05. **B)** Cellular localization of upregulated prey proteins identified using the Human Cell Map resource. **C)** Full heatmap of proteins upregulated in at least one bait. Data highlighted with a violet frame corresponds to the information presented in Figure 1E. **D)** Gene ontology analysis for the individual baits. **E)** Plotting of BioID data as log fold change (log FC) against -log10 (adjusted p-value, padj) for selected preys.

**Supplementary Figure 3.**
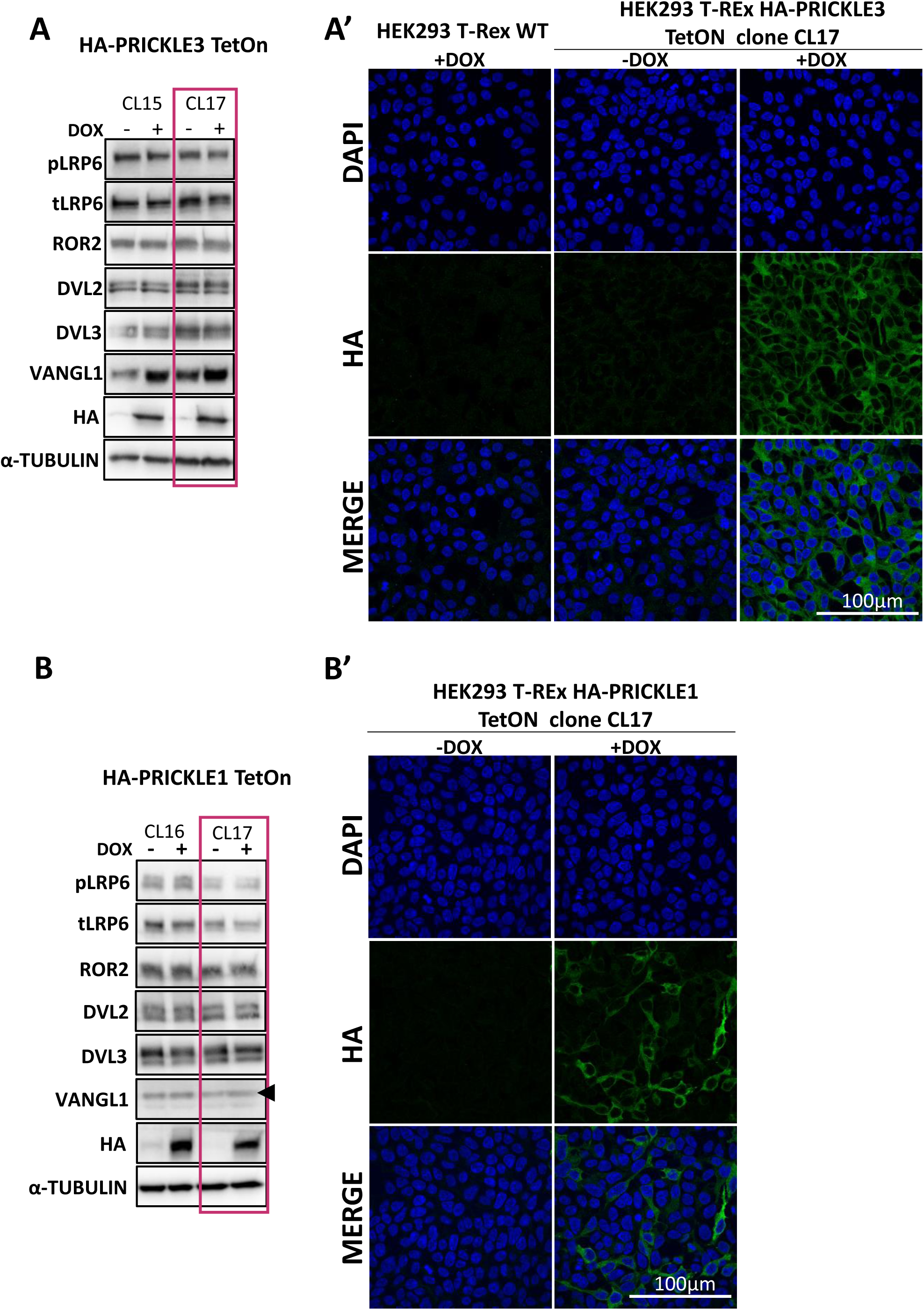
**A)** Validation of doxycycline-inducible cell lines expressing N-terminally HA-tagged PRICKLE3. The expression of PRICKLE3 and proteins representing components of both canonical and non-canonical WNT pathways was verified in clonally derived cells in the presence of doxycycline by Western blot. α-TUBULIN served as loading control. Clone 17 was selected for further research. **A’)** Immunofluorescence analysis of doxycycline-inducible cell lines expressing N-terminally HA-tagged PRICKLE3. Cells were stained with an anti-HA antibody to confirm the induction of PRICKLE3 expression. Nuclei were counterstained with DAPI. HEK T-REx 293 wild-type (WT) cells served as a specificity control for the anti-HA antibody. **B)** Validation of doxycycline-inducible cell lines expressing N-terminally HA-tagged PRICKLE1. The expression of PRICKLE1 and proteins representing components of both canonical and non-canonical WNT pathways was verified in clonally derived cells in the presence of doxycycline by Western blot. α-TUBULIN served as loading control. Clone 17 was selected for further investigation. **B’)** Immunofluorescence analysis of doxycycline-inducible cell lines expressing N-terminally HA-tagged PRICKLE1. Cells were stained with an anti-HA antibody to confirm the induction of PRICKLE1 expression. Nuclei were counterstained with DAPI.

**Supplementary Figure 4.**
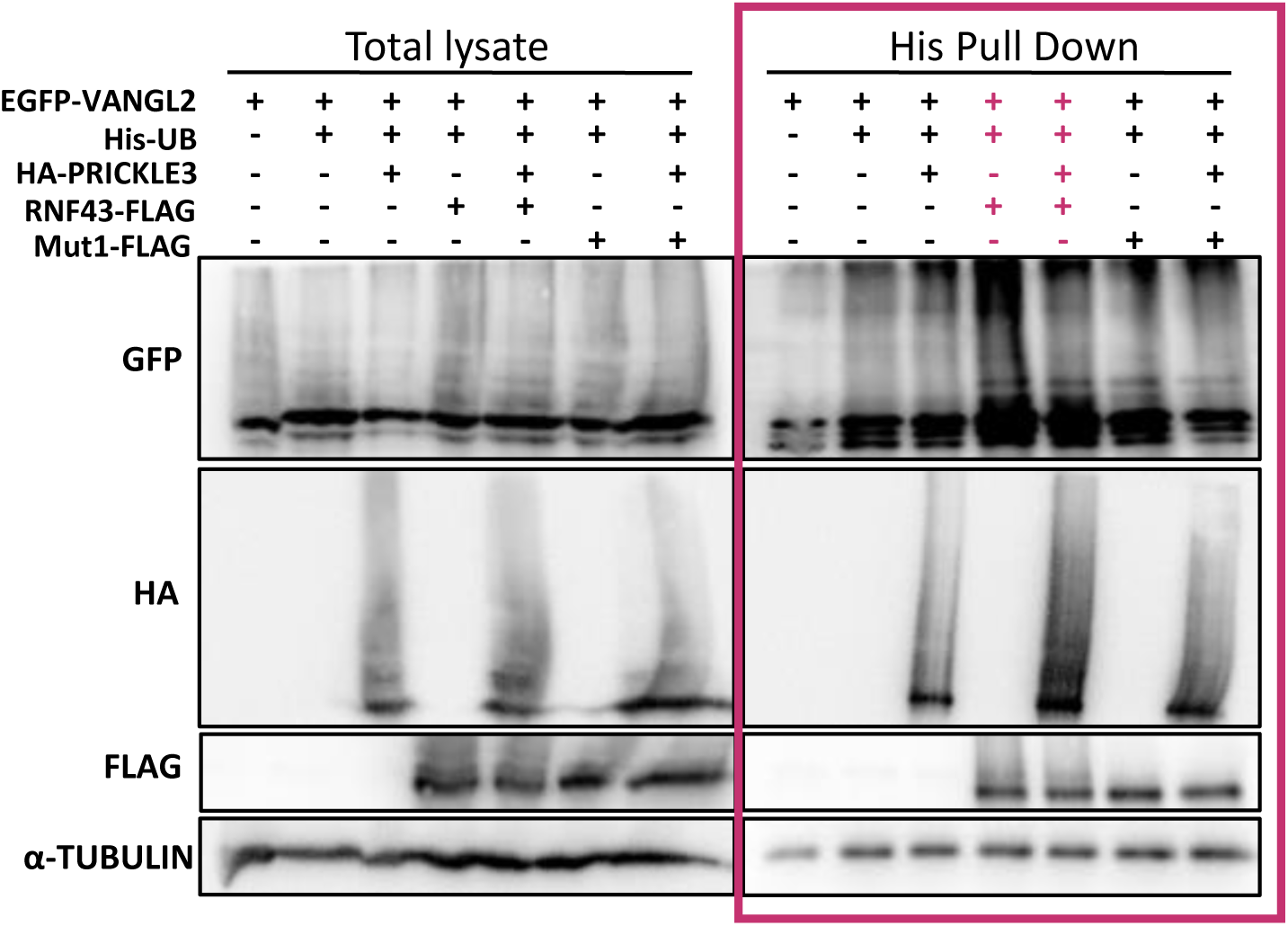
Ubiquitination assay. HEK T-REx 293 PRICKLE3 TetON cells were transfected His-tagged ubiquitin, EGFP-VANGL2 and FLAG-tagged wild-type or enzyme-dead Mut1 RNF43 constructs. PRICKLE3 expression was induced by ON treatment with doxycycline. Ubiquitinated proteins were enriched by His pull down and analysed by WB. Representative experiment from N = 3. Data highlighted by a violet frame corresponds to the information presented in Figure 1E.

**Supplementary Figure 5.**
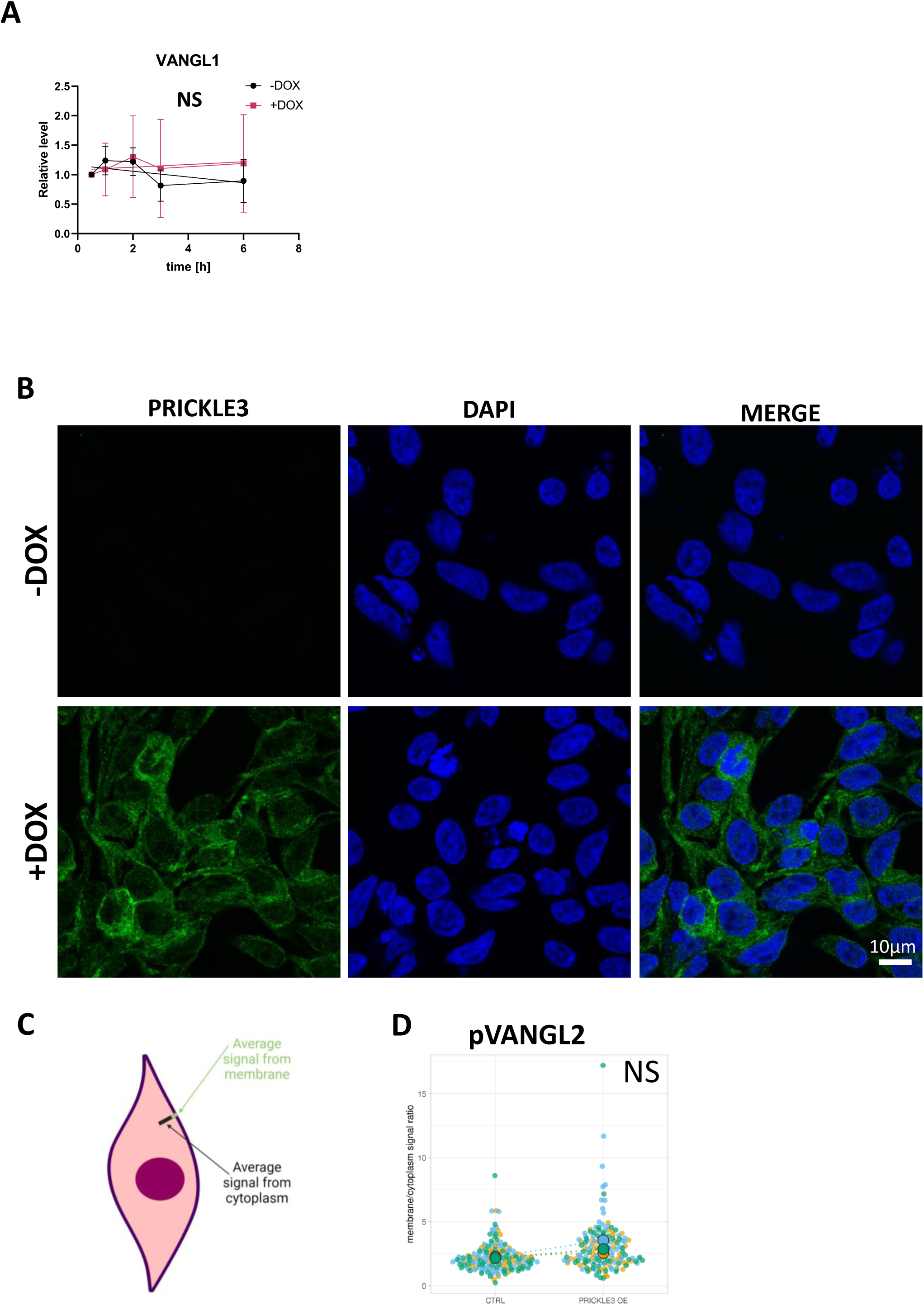
**A)** Densitometric quantification of Western blot (WB) band intensities. Results were normalized to the 0-hour time point. Statistical analysis was performed using linear regression. (NS) indicates no statistical difference. N = 5. Data corresponds to the information presented in Figure 5B. **B)** Immunofluorescence imaging of cell level of PRICKLE3. Cells were stained with an anti-HA antibody to confirm the induction of PRICKLE3 expression. Data corresponds to the information presented in Figure 5C. **C)** Schematic representation illustrating the quantification method for VANGL1 and pVANGL2 levels on the cell membrane. **D)** Quantification of the membrane levels of VANGL and pVANGL. Data corresponds to the information presented in Figures 5C and 5D. Statistical analysis was performed using the SuperPlots of Data tool, (NS) indicates no statistical difference. N= 3

## Supplementary Tables 1-13

See the Excel file

